# Learning probability distributions of sensory inputs with Monte Carlo Predictive Coding

**DOI:** 10.1101/2024.02.29.581455

**Authors:** Gaspard Oliviers, Rafal Bogacz, Alexander Meulemans

**Affiliations:** MRC Brain Network Dynamics Unit, Nuffield Department of Clinical Neurosciences, University of Oxford, Oxford, United Kingdom; Department of Computer Science, ETH Zurich, Zürich, Switzerland

## Abstract

It has been suggested that the brain employs probabilistic generative models to optimally interpret sensory information. This hypothesis has been formalised in distinct frameworks, focusing on explaining separate phenomena. On one hand, predictive coding theory proposed how the probabilistic models can be learned by networks of neurons employing local synaptic plasticity. On the other hand, neural sampling theories have demonstrated how stochastic dynamics enable neural circuits to represent the posterior distributions of latent states of the environment. Here, we bring together these two lines of theoretic work by introducing Monte Carlo predictive coding (MCPC). We demonstrate that the integration of predictive coding with neural sampling results in a neural network that learns precise generative models using local computation and plasticity. The neural dynamics of MCPC infer the posterior distributions of the latent states in the presence of sensory inputs, and can generate likely inputs in their absence. Furthermore, MCPC captures the experimental observations on the variability of neural activity during perceptual tasks. By combining predictive coding and neural sampling, MCPC offers a unifying theory of cortical computation which can account for both sets of neural data that previously had been explained by these individual frameworks.

## 1 Introduction

The Bayesian brain hypothesis states that the brain learns and updates probabilistic generative models of its sensory inputs. By learning accurate generative models, the brain establishes the causal relationship between environmental states and sensory inputs [1–3]. The brain also mitigates the effect of sensory noise through generative models by optimally integrating prior knowledge with new sensory data. Several studies have successfully employed probabilistic generative models to explain behavior [4–9], and interpret neural activity [10–14].

To elucidate how the brain represents generative models, we seek a neural network capable of learning generative models, while adhering to the brain’s intrinsic characteristics. These characteristics include (i) the brain’s ability to infer posterior distributions of environmental states given sensory inputs [4–7, 15], (ii) its proficiency in constructing accurate generative models using hierarchical neural networks [16], and (iii) its reliance on localized computation and plasticity within these networks [17, 18].

Multiple models implementing the Bayesian brain principle have been proposed that capture some of the above characteristics of the brain, however none adhere to all these constraints. Below we review two categories of models that focus on describing learning of probabilistic models and representing the posterior probabilities of the latent states respectively.

An influential theory describing how the cortex learns the generative models is predictive coding. It hypothesises that the brain learns the generative models by minimising the error between actual sensory inputs and the sensory inputs predicted by its model [19–21]. Predictive coding proposes a hierarchically structured neural network that is local in computation and plasticity. Moreover, predictive coding serves as a comprehensive framework for understanding attention [22], a range of neurological disorders [23], and various neural phenomena [19, 24, 25]. However, predictive coding has two limitations: First, it only infers the most likely state of an environment from sensory inputs, rather than the whole posterior distribution, thereby ignoring any uncertainty information [21]. Second, it demonstrated a limited learning performance for generative tasks [26]. Recent work extended predictive coding to improve its learning performance using lateral inhibition and sparse priors [26, 27], however the resulting neural network is still unable to infer posterior distributions. In addition to predictive coding, other models have been proposed to describe learning of probabilistic models in the brain. For example, a recent study has employed generative adversarial networks to explain delusions observed in some mental disorders [28]. However, no biologically plausible neural implementation of the adversarial objective function has been identified.

On the other hand, a wide range of neural sampling models have also been proposed that infer the posterior distributions using Monte Carlo sampling methods [12, 29–34]. In these models, the fluctuations of neural activity over time sample the probability distributions the brain is trying to infer. Some studies show that neural variability in the brain exhibits characteristics consistent with neural sampling processes [35–37]. Despite this, present neural sampling models lack learning capabilities, local learning rules, or depth in their neural architectures. Recent work incorporated neural sampling into a sparse coding model that can learn generative models with local plasticity [31]. However, the sparse coding model does not include a hierarchical architecture that can support learning of complex generative models.

Here, we bring together the above work on predictive coding and neural sampling by proposing Monte Carlo predictive coding (MCPC), a biologically plausible neural implementation of generative learning in the brain. Compared to previously published models of generative learning in the brain, MCPC is the first model to infer full posteriors and learn hierarchical generative models by relying solely on local computation and plasticity. MCPC is also able to generate sensory inputs using local neural dynamics, and its neural activity captures the variability in cortical activity during perceptual tasks. MCPC has robust learning capabilities across noise types and intensities as well. Through this integration, MCPC offers a comprehensive theoretical framework for understanding neural computation. Since we first presented MCPC at a conference [38], concurrent work in the field of machine learning [39, 40] showed that sampling in energy-based models, similar to MCPC, scales to challenging machine learning tasks with comparable performance to variational autoencoders [41]. These developments are complementary to our work on unifying neural sampling with predictive coding in biologically plausible networks capturing key characteristics of cortical activity.

## 2 Results

This section presents MCPC, and it is organized into subsections discussing the following properties of the model:

1. MCPC utilizes neural networks with local computation and plasticity to learn hierarchical generative models.
2. MCPC’s neural dynamics infer full posterior distributions of latent variables in the presence of sensory inputs.
3. MCPC’s neural dynamics sample from the learned generative model in the absence of sensory inputs.
4. MCPC learns accurate generative models of sensory data.
5. MCPC captures the variability of neural activity observed in perceptual experiments.
6. MCPC achieves robust learning capabilities across noise types and intensities.

Throughout our experiments, we consider two tasks: learning a simple probability distribution of Gaussian sensory data, and learning a more complex distribution of handwritten digit images from the MNIST dataset [42]. We compare the properties of our model to predictive coding (PC) [21], because MCPC builds upon PC, and the performance of PC has been characterised in a variety of tasks (it achieves performances similar to backpropagation in supervised machine learning tasks [43], and superior to backpropagation in tasks more similar to those faced by biological organisms [44]).

### 2.1 MCPC implementation with local computation and plasticity

To describe MCPC, we will first define a hierarchical generative model MCPC assumes, next present its inference and learning algorithm, and then show how it can be implemented through local computation and plasticity.

MCPC learns a hierarchical Gaussian model of sensory input *y* with latent variables *x*. The latent variables are organized into *L* layers in this model. We denote the activity of sensory neurons by *x*_0_, and when the sensory input is present, they are fixed to it, i.e., *x*_0_ = *y*. Sensory input *y* is predicted by the first layer *x*_1_ while variables *x*_*l*_ in layer *l* are predicted by the layer above. The resulting joint distribution over sensory inputs and latent variables is given by:

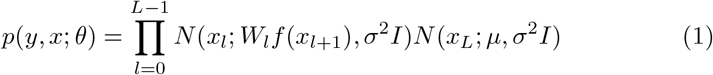

where *x* denotes the latent states *x*_1_ to *x*_*L*_, parameters *θ* comprise weights *W*_*l*_ and the prior mean *µ* describing the mean activity in the top layer, *f* stands for an activation function, *I* represents an identity matrix, and *σ*^2^ denotes a scalar variance. A simple example of such probabilistic model is illustrated in figure 1a, and it includes one sensory input and one latent state. Such model could for instance be used by an organism to infer the size of a food item based on observed light intensity [21]. We will use this model throughout the paper to provide intuition before considering more complex models.

**Figure 1.**
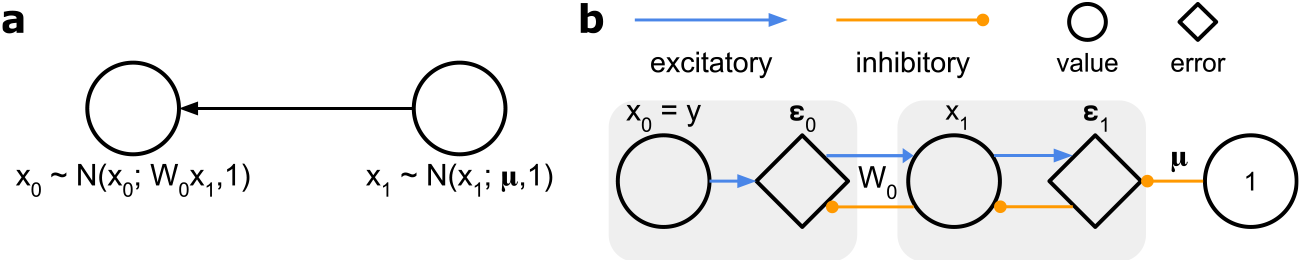
Example of a probabilistic model and its corresponding neural implementation for MCPC. **a**, Linear Gaussian model with one sensory input and one latent state. **b**, The neural implementation of MCPC using local synaptic connections for this model.

MCPC learns a hierarchical Gaussian model by iterating over two steps that descend the negative joint log-likelihood

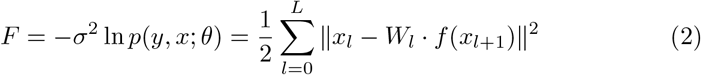

In the first step, MCPC leverages Markov chain Monte Carlo techniques to approximate the full posterior distribution by using the following Langevin dynamics [45]:

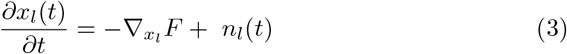

Thus we modify the latent variables to reduce *F*, but additionally add a zero-mean noise *n*_*l*_(*t*) (in the next subsection we will show explicitly that such dynamics lead to *x*_*l*_ sampling from its posterior distribution). The noise needs to be uncorrelated over time, i.e. with covariance 𝔼 *n*_*l*_(*t*)*n*_*l*_(*t*^*′*^)^⊤^ = 2*σ*^2^*δ*(*t* − *t*^*′*^)*I*, where *δ* is the Dirac delta function and *I* the identity matrix. Here, the noise variance *σ*^2^ equals the layer variance of MCPC’s hierarchical Gaussian model (equation (1)). Note that the layer variance of the model *σ*^2^ can also be encoded in the joint log-likelihood *F*, as is common in predictive coding [21, 46] (see Supplementary information 1). However, we chose to encode layer variance in the noise variable to make the dependence between the level of noise in MCPC’s dynamics and the variance of its generative layers explicit.

Evaluating the gradient in equation (3), we see below that these neural dynamics give rise to prediction errors *ϵ*_*l*_ encoding the mismatch between the predicted latent state *W*_*l*_*f* (*x*_*l*+1_) and the inferred latent state *x*_*l*_.

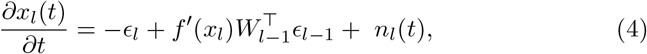

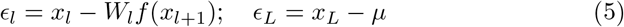

In the second step, MCPC uses the noisy neural activities to update its parameters as follows:

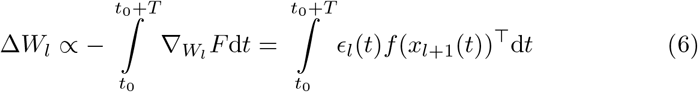

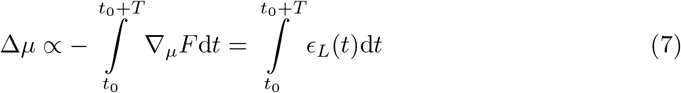

with *t*_0_ the time point where the noisy dynamics have converged to their steadystate distribution, and *T* is large to ensure that the dynamics sample from the whole steady-state distribution. Repeating these two steps enables inference of latent variables and learning of model parameters.

The above algorithm has a direct implementation in a neural network. Such a network has two classes of neurons: value neurons encoding latent states and error neurons encoding prediction errors. The weights of synaptic connections in such a network encode the parameters of the generative model. This is illustrated in figure 1b through a simple network implementing probabilistic inference in the model from figure 1a.

The neural network of MCPC relies on local computation. All neurons perform computations solely based on the activity of their input neurons and the synaptic weights related to these inputs. Specifically, the rate of change of value neurons in equation (4) depends on their own activity, the activity of the error neurons connected with them, the weights of these connections, and local noise.

Similarly, the activity of error neurons in equation (5) can be computed using the activity of connected value neurons and corresponding synaptic weights.

The network also exhibits local plasticity. Synaptic plasticity in MCPC (equations (6-7)) relies exclusively on the product of the activity of pre-synaptic and post-synaptic neurons. The integral in MCPC’s synaptic plasticity can also be approximated using local plasticity. This could be achieved by continuously updating synaptic weights with a large time constant.

The neural dynamics and parameters updates of MCPC prescribe the same local neural circuits as existing implementations of predictive coding [21, 47], with the addition of a noise term. Hence, MCPC shares the focus of predictive coding on minimizing prediction errors. The additional noise term does, however, lead to significant benefits as discussed below.

### 2.2 MCPC infers posterior distributions

Here we show that MCPC’s neural activity infers full posterior distributions of latent variables in the presence of sensory inputs. We prove that MCPC’s neural activity samples the posterior *p*(*x*|*y*; *θ*) at its steady state for an input *y*. Moreover, we confirm that MCPC’s neural activity approximates the posterior in the linear model of figure 1a and in a model trained on MNIST digits.

Proposition 1 demonstrates that the neural activity *x* prescribed by MCPC samples from the posterior *p*(*x*|*y*; *θ*) over latent states *x* when the dynamics in equation (3) have converged.

#### Proposition 1

*The posterior p*(*x*|*y*; *θ*) *is the steady-state distribution p*^*ss*^(*x*) *of the inference dynamics of MCPC:*

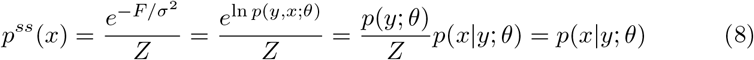

*where Z is the partition function*.

The proof is given in the transformations in equation (8), which we now explain. It follows from a classical result in statistical physics that the steadystate distribution *p*^*ss*^(*x*) of a variable *x* described by the Langevin equation

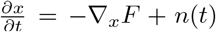 is given by 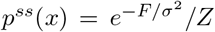 when the variance of the noise *n*(*t*) equals 2*σ*^2^ [48]. The Langevin dynamics of MCPC minimise the joint log-likelihood *F* = − *σ*^2^ ln *p*(*y, x*; *θ*). The distribution *p*^*ss*^(*x*) can therefore be rewritten as *p*(*y, x*; *θ*)*/Z*. Employing the conditional probability formula allows *p*^*ss*^(*x*) to be subsequently expressed as *p*(*y*; *θ*)*p*(*x*|*y*; *θ*)*/Z*. Given that the distribution *p*(*y*; *θ*) remains constant for a particular stimulus *y*, 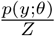 forms the partition function of the posterior *p*(*x*|*y*; *θ*). However, the posterior distribution integrates to one, ∫ *p*(*x*|*y*; *θ*) *dx* = 1. This implies that 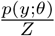 equals one and that the steady-state distribution *p*^*ss*^(*x*) effectively simplifies to *p*(*x*|*y*; *θ*).

To verify this property, we validate that MCPC samples from the posterior distribution within the simple model from figure 1a, which is tractable. Figure 2a illustrates the activity of latent state *x*_1_ of this model during inference under a constant input for both MCPC and PC. While the activity converges to a single value for PC, activity for MCPC fluctuates around this value representing the uncertainty in its inference. Figure 2b displays a histogram of the latent state’s activity over time throughout the MCPC inference. The inference of PC at its convergence point is also illustrated, as well as the posterior distribution *p*(*x*_1_ |*y*; *θ*) for the specified input. MCPC’s latent state activity accurately samples the posterior of the linear model. In contrast, PC’s inference converges to the mode of the posterior. This result confirms that MCPC samples from the posterior *p*(*x* | *y*; *θ*), whereas PC infers the Maximum a-posteriori (MAP) estimate.

**Figure 2.**
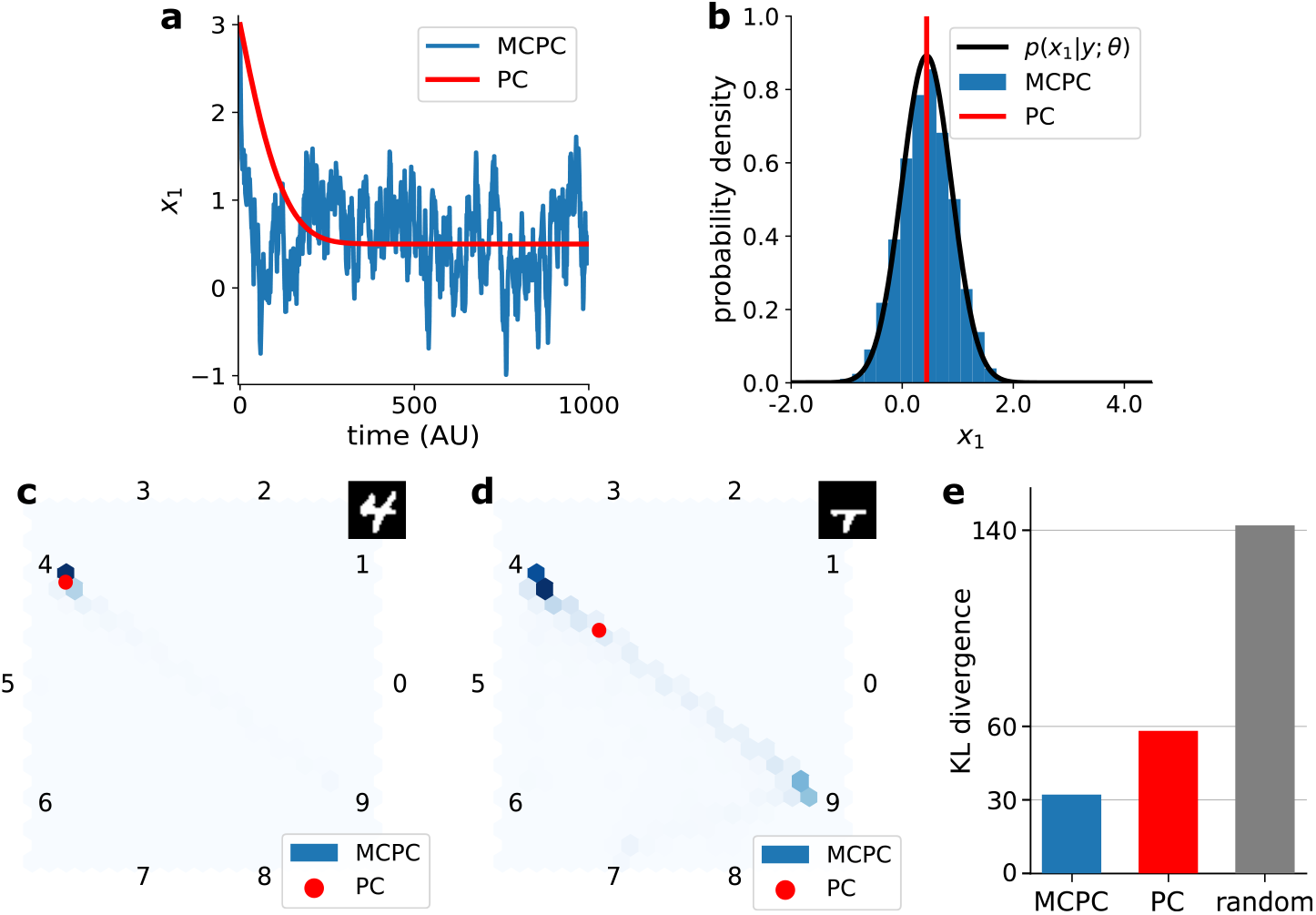
Neural activity of MCPC infers posterior distributions in the presence of inputs. **a**,**b**, Latent state activity *x*_1_ of MCPC and PC in the linear model shown in figure 1a with parameters {*W*_0_ = 2, *µ* = 0.5} and input *y* = 1. **c**,**d**, Latent state activity of MCPC and PC in a model trained on MNIST with a digit image and a half-masked digit image (see top-right) as input. Plots (**b**), (**c**), and (**d**) show a histogram of MCPC’s activity over 10,000 timesteps and PC’s activity at converges. **e**, KL divergence between the digit class distribution inferred by an ideal ResNet-9 observer and the class distribution decoded from the latent state *x*_*L*_ inferred by MCPC and PC for masked digit images. The KL divergence for shuffled distributions is also provided. Animation of the MCPC’s latent activity in plots b to d can be found in Supplementary videos 1-3.

Next, we visually confirm that MCPC infers latent states correctly in a non-linear model with three latent layers trained on MNIST digits. Visualising the latent states during inference shows that both MCPC and PC infer the correct digit when provided with a full-digit image (Figure 2c). However, when prompted with an ambiguous masked-digit image, MCPC identifies different possible interpretations, while PC only infers one possible interpretation (Figure 2d). This result indicates that MCPC approximates the posterior distribution more accurately than PC. The visualisations are obtained by employing a linear classifier to interpret the latent states. This classifier decodes the latent state *x*_*L*_ and generates a probability distribution over the ten-digit categories. This distribution can then be visualised by mapping it onto ten evenly spaced unit vectors within a circle [49](see Methods section 4.2.2 for details).

Finally, we show quantitatively that MCPC indeed approximates the posterior better than a MAP estimate in a non-linear model trained on MNIST. Figure 2e shows the Kullback–Leibler (KL) divergence between the posterior across digit classes inferred by a ResNet-9-based ideal observer and the distributions inferred by MCPC, and PC for half-masked images. The KL divergence for a random baseline obtained with shuffled distributions is also shown. This figure shows that the KL divergence between the distributions inferred by the ideal observer and by MCPC is smaller than the one for PC and for the baseline. The lower KL divergence confirms that MCPC’s inferred latent states capture the posterior distribution more accurately than PC’s MAP estimate. In this experiment, ResNet-9 is a classifier that achieves over 99% classification accuracy on MNIST [50]. The probability distributions across digit classes of MCPC and PC inferences are obtained with the linear classifier used for interpreting the latent states. Moreover, the random baseline is calculated by averaging the KL divergence between the inferences of the ideal observer and the shuffled distributions inferred by MCPC and by PC.

### 2.3 MCPC samples from its generative model in the absence of sensory inputs

Here we show that in the absence of sensory inputs, MCPC spontaneously samples from its learned generative model of sensory inputs. We prove that the activity of the unclamped input neurons sample from probability distributions of sensory inputs learned by MCPC. Experiments confirm that the unclamped neural activity generates sensory inputs learned by MCPC in the simple model of figure 1a and in a model trained on MNIST.

To model a scenario in which no sensory input is provided, instead of clamping the input neurons *x*_0_ to a sensory stimulus *y*, we let these neurons follow similar Langevin dynamics as all other neurons:

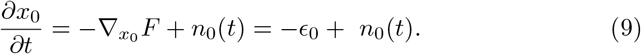

Proposition 2 shows that when input neurons are not fixed to sensory stimuli, MCPC spontaneously samples from the learned probability distribution of sensory inputs. In this proposition, we demonstrate that the steady state of MCPC’s unclamped activity is equal to the marginal likelihood *p*(*x*_0_; *θ*).

#### Proposition 2

*The marginal likelihood p*(*x*_0_; *θ*) *is the steady-state distribution p*^*ss*^(*x*_0_) *of the Langevin dynamics given in equation (9):*

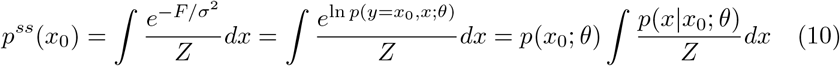

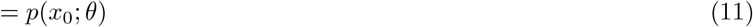

The proof of proposition 2 is similar to that of proposition 1. The steadystate distributions of the neural activity in MCPC in the absence of an input *p*^*ss*^(*x*_0_, *x*) is given by 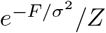. This is a consequence of the Langevin dynamics of MCPC minimizing the joint log-likelihood *F* while subjected to a noise variable with variance 2*σ*^2^. This distribution can be marginalised over the latent states *x* = [*x*_1_, …, *x*_*L*_] to find the steady-state distribution of the sensory input neurons *p*^*ss*^(*x*_0_). The joint log-likelihood *F* equals − *σ*^2^ ln *p*(*y, x*; *θ*), where *y* = *x*_0_ when input neurons are unclamped. The expression for *p*^*ss*^(*x*_0_) is therefore reformulated as 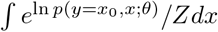. This expression can be rewritten as ∫*p*(*x*_0_) *p*(*x*|*x*_0_)*/Zdx*. Given that the expression ∫*p*(*x*|*x*_0_)*/Zdx* remains constant for a specific activity *x*_0_, this expression forms the partition function of the marginal likelihood *p*(*x*_0_; *θ*). However, the marginal likelihood integrates to one, ∫*p*(*x*_0_; *θ*) *dx* = 1. This implies that the partition function ∫*p*(*x* |*x*_0_)*/Zdx* equals one and that the steady-state distribution *p*^*ss*^(*x*_0_) effectively simplifies to *p*(*x*_0_; *θ*).

Figures 3a and 3b experimentally confirm that MCPC generates accurate samples of the generative distribution in the absence of sensory inputs. Figure 3a demonstrates this for the linear model by showing that the activity of the unclamped input neuron matches the model’s generative distribution *p*(*x*_0_; *θ*). Similarly, figure 3b illustrates that the unclamped input neurons of a deep non-linear model trained on MNIST produce activity patterns that resemble the digit images used in training.

**Figure 3.**
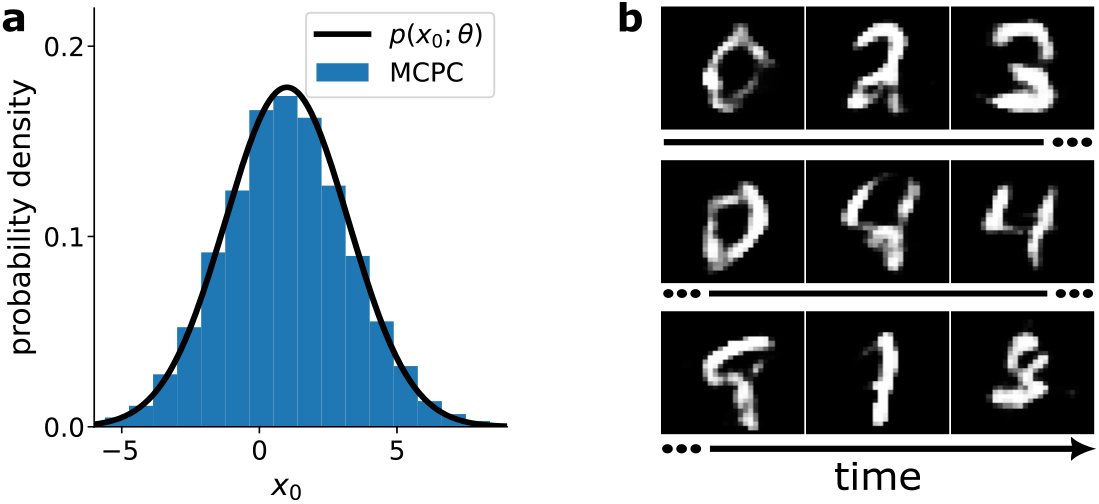
Neural activity of MCPC samples its generative model in the absence of inputs. **a**, Histogram of the MCPC activity of the unclamped input neuron *x*_0_ in the linear model given in figure 1a with parameters *{W*_0_ = 2, *µ* = 0.5*}* obtained over 10,000 timesteps. **b**, Activity of input neurons *x*_0_ in a model trained on MNIST displayed for time points separated by 3,000 timesteps. Animation of the MCPC’s unclamped input activity can be found in Supplementary videos 4 and 5.

### 2.4 MCPC learns accurate generative models

We show here that MCPC learns precise generative models of sensory data. We demonstrate the precise learning of MCPC by first proving that MCPC is guaranteed to learn locally optimal generative models. Afterwards, we experimentally confirm that MCPC learns accurate generative models of Gaussian sensory data and handwritten digit images. In the process, MCPC outperforms PC and approaches the performance of Deep Latent Gaussian models (DLGMs) on the digit learning task. DLGMs are the standard machine learning approach for training hierarchical Gaussian models (equation 1) using backpropagation [51], and are therefore used as a benchmark.

MCPC learns locally optimal generative models of sensory data by implementing the Monte Carlo expectation-maximization algorithm (see proposition 3). This algorithm guarantees convergence to locally optimal parameters when given enough sampling time during inference [52].

#### Proposition 3

*MCPC implements the Monte Carlo expectation-maximization algorithm by iterating over:*

1. *E-step: MCPC’s inference, x*(*t*), *approximates the posterior distributions using an MCMC method for a given input y* 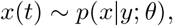
2. *M-step: MCPC’s parameter update maximizes the Monte Carlo expectation of joint log-likelihood* 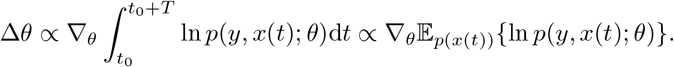

Proposition 3 relies on proving that MCPC’s inference samples the posterior distribution for a given input and proving that MCPC’s parameter updates maximize the Monte Carlo expectation of the joint log-likelihood. Proposition 1 shows that MCPC’s inference samples the posterior distribution. This provides half of the proof for proposition 3. The second part of the proof can be shown by identifying that the expressions *F* in equations (6) and (7) equal the scaled joint log-likelihood. This allows MCPC’s parameter updates to be rewritten as 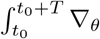 ln *p*(*y, x*(*t*); *θ*)d*t*. The partial derivatives can be taken out of the integrals to obtain the parameter update 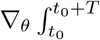. In this expression, 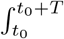In. *p*(*y,x*(*t*);θ)*dt* is the Monte Carlo expectation of the joint log-likelihood. Consequently, MCPC parameter updates maximise the Monte Carlo expectation of joint log-likelihood.

Additionally, we show that MCPC accurately learns the distribution of Gaussian sensory data with the linear model of figure 1a. Figure 4a illustrates the distribution learned by MCPC after 375 parameter updates. This distribution accurately models the Gaussian data distribution used for training. We obtain the samples of the distribution learned by MCPC using ancestral sampling. In a hierarchical Gaussian model, ancestral sampling consists of first sampling the top latent layer *x*_*L*_ from its Gaussian distribution 𝒩 (*x*_*L*_; *µ I*). Each layer is then sampled sequentially using the conditional Gaussian distribution 𝒩 (*x*_*l*_; *W*_*l*_*f* (*x*_*l*+1_), *I*). Figure 4b verifies that MCPC learns an accurate model of Gaussian data for different model initialisations. This figure demonstrates that each parameter trajectory converges to the parameters for ideal data modeling. This convergence can also be validated analytically by first calculating the curves where the parameter update for the weight or the prior mean parameter equals zero (these curves are known as nullclines and shown in green and purple in figure 4b). The intersection of these curves provides the equilibrium points for the parameter values. For MCPC, this intersection is located at the model parameters that perfectly capture the Gaussian data distribution (see Supplementary information 2 for full derivation).

**Figure 4.**
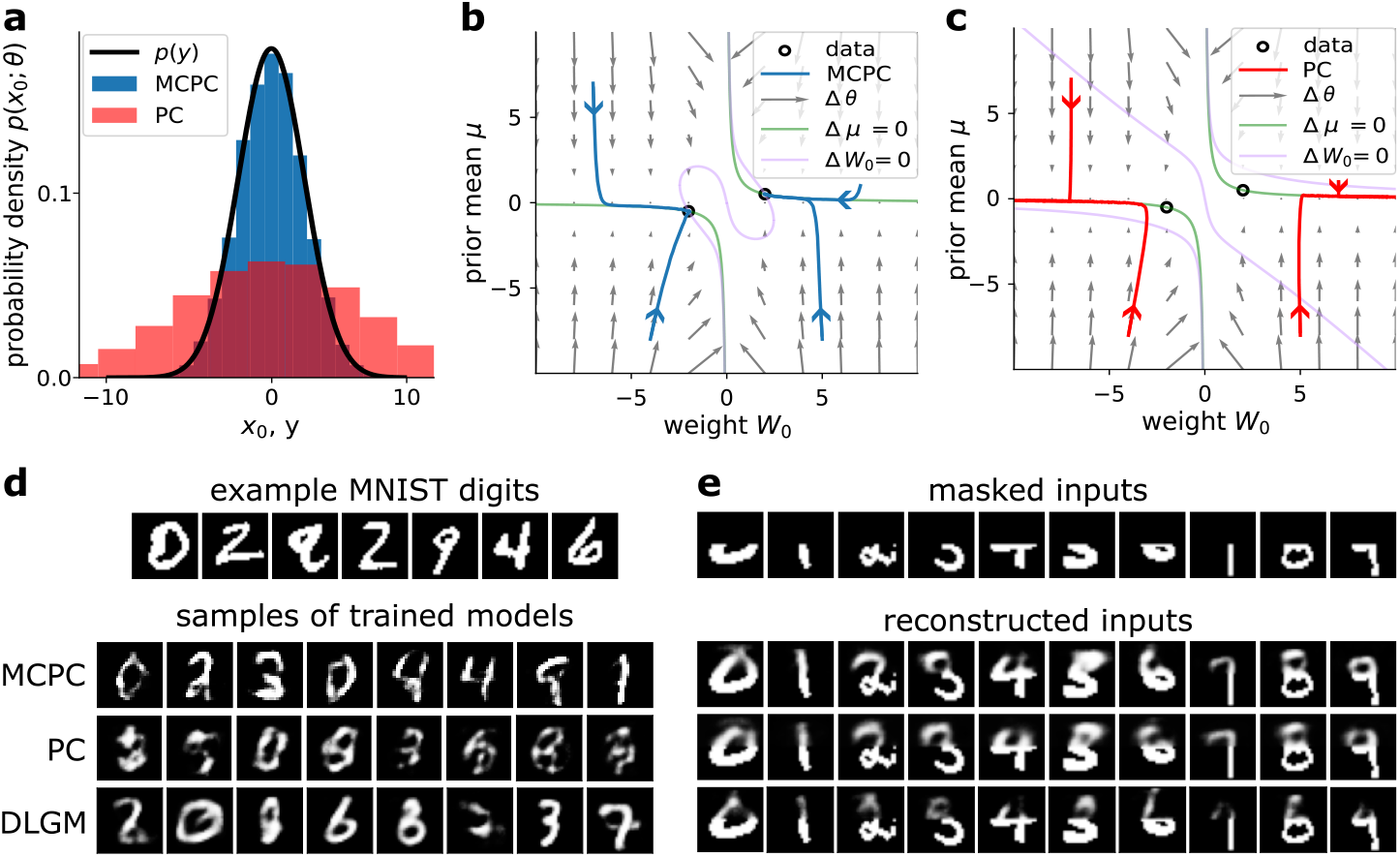
MCPC learns accurate generative models of sensory inputs. **a**, Distributions learned by MCPC and PC in the linear model given in figure 1a after 375 parameter updates. **b**,**c**, Evolution of the weight *W*_0_ and prior mean *µ* parameter of the linear model during training with MCPC (**b**) and PC (**c**). The optimal model parameter values are marked as hollow dots. The vector field shows the expected gradient flow of the parameters. The additional curves reveal nullclines where the parameter update for the weight or the prior mean parameter equals zero (see Supplementary information 2 for derivations). **d**, Comparison between samples obtained from models trained with MCPC and PC on MNIST, as well as from a DLGM trained on MNIST. The samples are obtained by ancestrally sampling the models for PC and the DLGM and by sampling the spontaneous neural activity for MCPC. **e**, Comparison between masked images reconstructed by MCPC, PC, and a DLGM. We reconstruct the images by obtaining a Maximum a-posteriori estimate of the missing pixel values.

In contrast to MCPC, PC learns a strikingly poor generative model of the Gaussian data as shown in figure 4a. PC learns a Gaussian distribution with the correct mean but with an excessive variance. This high variance is caused by the diverges of PC’s weight to *±*∞ during training as shown in figure 4c for different model initialisations. The variance of the model learned by PC equals 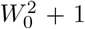 (see equation (19) in Methods). Consequently, the variance of PC’s generative distribution grows toward infinity as training progresses, leading to a model that becomes increasingly inaccurate. PC’s parameter *W*_0_ diverges to*±*∞ additionally validating the suboptimal learning performance of PC (see Supplementary information 2). The underlying cause of this undesirable behavior of PC is that it uses the maximum a-posteriori estimate of *x*_*l*_ in its parameter updates, instead of the full posterior distribution. This learning strategy is equivalent to variational expectation maximization [53], with a Diracdelta as variational approximation to the true posterior. Crucially, the Diracdelta introduces an infinite entropy, causing PC to optimize a bound on ln *p*(*y*; *θ*) which is arbitrarily loose (refer to Supplementary information 3 for additional details). In contrast, MCPC optimizes a tight bound on ln *p*(*y*; *θ*) (Proposition 3). Interestingly, a range of other theories for learning in the brain [43, 54, 55] are based on a similar energy as in PC, posing the question of whether they suffer from similar failure modes as we uncover here for PC.

Next, we show that MCPC learns accurate hierarchical Gaussian models of MNIST handwritten digit images [42]. For this learning task, we consider non-linear models with three latent layers. We train these models using MCPC, PC, and the DLGM approach (refer to methods section 4.2.2 for details). Figure 4d presents samples generated from the trained models. The quality of samples generated from an MCPC-trained model approaches that of samples produced by a DLGM. However, the samples obtained from training with PC are of significantly poorer quality, even when we apply weight decay to mitigate PC’s exploding variance. To quantify the difference in performance, we compute three metrics. First, we calculate the Fréchet inception distance (FID) of the generated samples which measures the similarity between generated data and actual data [56]. Second, we approximate the marginal log-likelihood of test data ln *p*(*y*_*eval*_; *θ*) using Monte Carlo sampling. This metric evaluates the generalization performance of a trained generative model. Finally, we compute the mean squared error (MSE) associated with reconstructing masked digits as illustrated in figure 4e. This assesses the ability to learn and retrieve associative memories [57]. Table 1 summarises the results and shows that a model trained with MCPC generates significantly better samples than PC and that it has better generalization performance. Additionally, MCPC approaches the generative learning performance of DLGM. Table 1 also shows that MCPC can reconstruct masked digits as well as PC and that both perform significantly better than DLGMs.

**Table 1.** Comparison of learning performance between MCPC, PC and DLGM. We report the FID, the marginal log-likelihood, and the reconstruction error on an MNIST evaluation set (the closer to zero the better for all metrics). We set in bold the best score across the models. Mean *±* standard deviation computed over three seeds.

| Model | FID | $-\ln p(y_{eval})$ | Reconstruction MSE ($10^{-2}$) |
| --- | --- | --- | --- |
| PC | $115.2 \pm 3.0$ | $168.9 \pm 0.2$ | $8.73 \pm 0.03$ |
| MCPC | $60.6 \pm 2.9$ | $144.6 \pm 0.7$ | **$8.29 \pm 0.05$** |
| DLGM | **$45.4 \pm 0.7$** | **$126.0 \pm 0.3$** | $12.04 \pm 0.08$ |

### 2.5 MCPC captures the variability of cortical activity

MCPC captures the key characteristics of the variability of cortical activity during perceptual tasks that PC fails to capture. Specifically, MCPC accounts for the suppression of neural variability at stimulus onset and the increase in similarity between spontaneous and evoked neural activities during development.

MCPC exhibits a decrease in temporal variability of neural activity at stimulus onset as observed in multiple electrophysiology studies [58–65]. These studies have shown that neural variability is smaller after stimulus onset than before stimulus onset. This finding holds when measured with intracellular or extracellular recordings and when an animal is task-engaged, awake, or anesthetized. Figure 5a illustrates the neural variability experimentally observed by Churchland et al. [61] and the neural variability of MCPC’s latent states at stimulus onset. This figure shows that MCPC’s neural activity captures the decrease in neural variability at stimulus onset for an MNIST-trained model. Supplementary information 4.1 provides additional proof that this observation generally holds for MCPC. This proof shows that the variability of MCPC’s steady state activity before stimulus onset is in expectation larger than the variability after stimulus onset.

**Figure 5.**
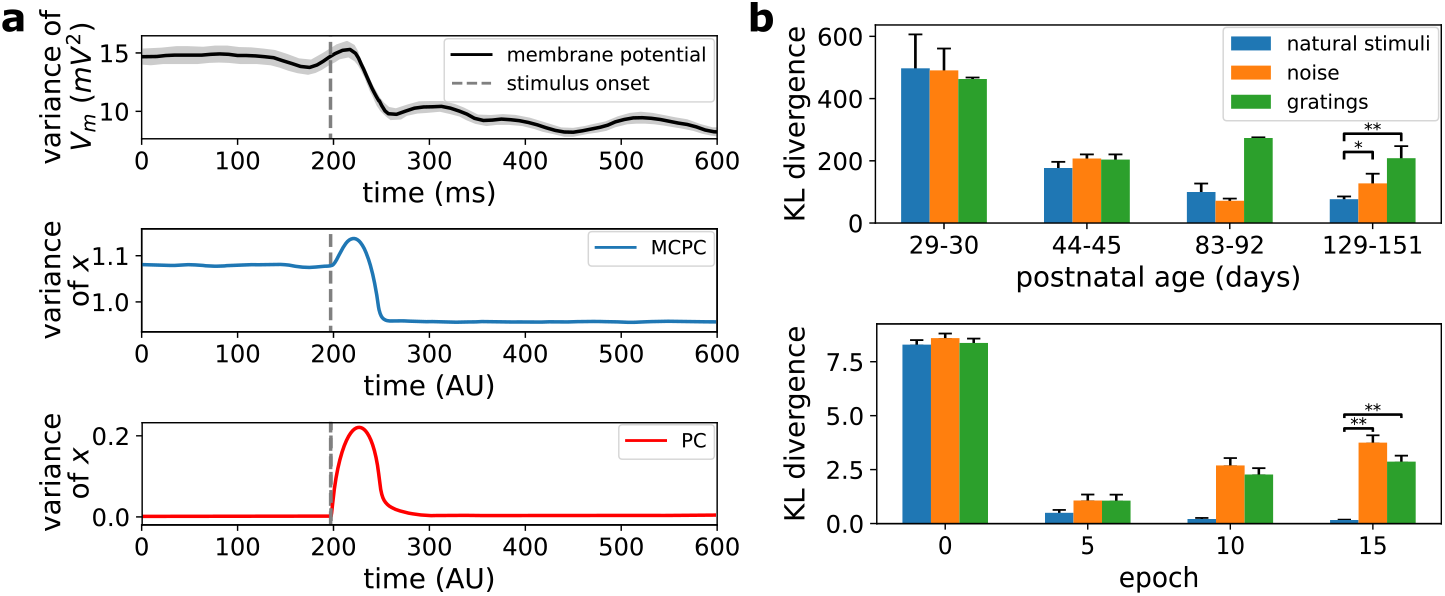
MCPC captures two key features of cortical activity. **a**, MCPC displays the decrease in neural variability at stimulus onset observed in the primary visual cortex (V1) of cats. The top plot recreates the neural quenching observed in the cortex (data re-plotted from figure 2c in Churchland et al. [61]). The middle and bottom plots show the mean temporal variability at stimulus onset of the latent state for MCPC and PC in a model trained on MNIST. Shaded regions give the s.e.m. Note that for MCPC and PC, these shaded regions are not visible due to their minimal magnitude. **b**, MCPC displays the similarity increase between spontaneous and evoked neural activities specific to natural stimuli observed in V1 of ferrets during development. The similarity is measured using the KL divergence between the distribution of spontaneous activity and the average distribution of evoked neural activities (the closer to zero the more similar). The average distribution is obtained for natural stimuli, noise stimuli, and gratings. The top plot recreates the similarity increase observed in awake ferrets (data re-plotted from figure 4a in Berkes et al. [35]). The bottom plot demonstrates a parallel increase in similarity specific to the training stimuli for MCPC. The MCPC model was trained on MNIST and evaluated using noise, image gratings, and MNIST digits (analogous to natural stimuli). In both plots, the error bars give the s.e.m. and * or ** indicate *p <* 0.05 or *p <* 0.01 respectively for a one-tailed paired samples t-test based on the KL divergences obtained for *n* = 10 MCPC models with the same architecture but different initializations.

MCPC displays an increase in similarity between spontaneous and average evoked neural activities that is specific to natural scenes as observed during learning for ferrets [35]. Berkes et al. [35] recorded the spontaneous and average evoked neural activity in V1 of ferrets for natural stimuli, sinusoidal gratings, and random noise. They observed that, as development progressed, the spontaneous activity increasingly resembled the average activity evoked by natural stimuli. Additionally, this increase in similarity was not observed for the sinusoidal gratings and random noise. Figure 5b compares the similarity between spontaneous and evoked neural activities for natural stimuli, noise, and image gratings reported by Berkes et al. [35] and observed for MCPC in MNIST-trained models. MCPC displays an increase in similarity between spontaneous and evoked neural activities that is specific to the digit stimuli on which it was trained (that are analogous to natural scenes to which the visual systems of animals were exposed). Such an increase in the similarity between spontaneous activity and the average response to stimuli on which the model was trained holds for MCPC in general. This is because the steady-state distribution of MCPC’s spontaneous neural activity becomes more similar to the average steady-state distribution of MCPC’s evoked activity as MCPC’s generative model improves (see Supplementary information 4.2 for proof). In our experiment, the similarity in neural activities is measured using the KL divergence and natural images are MNIST images (see Methods section 4.2.2 for a detailed explanation of the experiment).

Both the above characteristics of cortical activity are not reproduced by PC. The spontaneous and evoked neural activities of PC converge to constant neural activity without neural variability. Consequently, the temporal variability of individual neurons is not suppressed at stimulus onset in PC. Instead, the variability temporarily increases above zero at stimulus onset after which it returns to zero as illustrated in figure 5a. Moreover, training does not enhance the similarity between the distributions of spontaneous and evoked activities in PC. The distribution of PC’s spontaneous activity is a Dirac delta distribution as the activity has no variability. Similarly, the distribution of PC’s average evoked activity is a Dirac mixture distribution. Consequently, the KL divergence between these two non-identical Dirac-based distributions is always infinite.

### 2.6 MCPC learns effectively across noise types and intensities

A noteworthy characteristic of the MCPC is its flexibility in accommodating any type of noise distribution and variance, thereby avoiding the introduction of biologically unrealistic assumptions in the model. The only requirements on the noise variable *n*_*l*_(*t*) in MCPC’s dynamics are as follows: (1) the noise has a zero mean, (2) it is uncorrelated in time and across neurons, and (3) the variance of the noise needs to be constant over time. These requirements follow from the fluctuation-dissipation theorem in statistical mechanics that determines the first two moments of *n*_*l*_(*t*) (equations 12a-12b) and the resulting steady-state distribution (equation 13) where *σ*^2^ scales the variance of the noise [48].

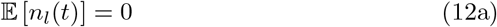

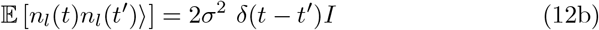

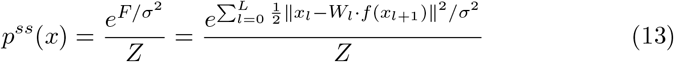

The fluctuation-dissipation theorem does not impose any specific constraints on the exact distribution of MCPC’s noise. This is due to the accumulation of uncorrelated noise at every instant in continuous time, ultimately leading to a Gaussian outcome at any non-zero time interval as a consequence of the central limit theorem [66]. This absence of assumption regarding the specific noise distribution ensures that MCPC does not hinge on potentially biologically implausible noise distributions.

The scalar variance of the noise, *σ*^2^, is not specified in the requirements of the fluctuation-dissipation theorem either. As a result, it can be arbitrarily assigned. However, it needs to be constant over time to guarantee consistency during learning and subsequent inferences. Moreover, the variance of the Gaussian input layers, which equals *σ*^2^*I* (see equation (13)), must remain lower than the variance of the data distribution to ensure accurate modeling of the data. To verify this result, we train the linear model from figure 1a with various levels of noise on Gaussian data.

We, then, compare the generative distribution and weight parameter *W*_0_ learned by MCPC to the data distribution and the ideal value of *W*_0_ respectively. The generative distributions are obtained by MCPC’s neural activity in the absence of inputs while maintaining the level of noise used during training. Figure 6a illustrates the generative distributions learned by MCPC, confirming that MCPC learns accurate generative models using constant noise with small variance *σ*^2^. Figure 6b reinforces this result by comparing the variance of the data distribution to the variance of the distribution learned by MCPC for a range of noise levels. MCPC learns a generative model with the correct variance, for *σ*^2^ smaller than the variance of the data. Finally, figure 6c demonstrates that MCPC successfully determines the optimal model weight *W*_0_ under varying noise levels. When the input layer variance *σ*^2^ surpasses the data variance, an optimal parameter set does not exist. In such cases, MCPC maximally reduces the model’s variance by setting the weight *W*_0_ close to zero.

**Figure 6.**
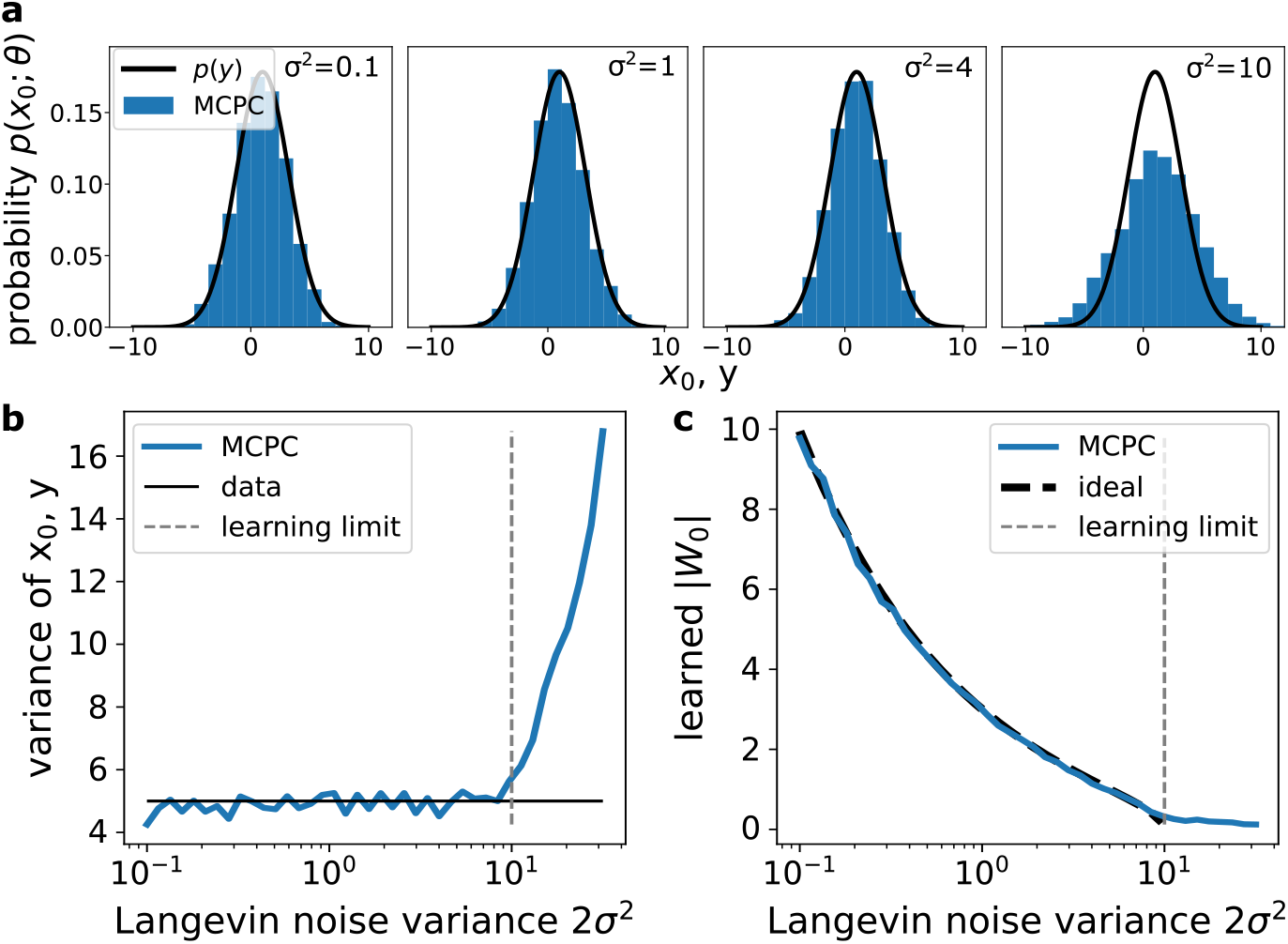
MCPC is compatible with a range of noise levels. **a**, Distributions learned by MCPC for the linear model shown in figure 1a of the main text when trained on Gaussian data with four different levels of noise. **b**, Comparison between the variance of the data distribution and the variance of the distribution learned by MCPC with a range of noise levels. **c**, Comparison between the weight parameter *W*_0_ learned by MCPC and the ideal weight for different levels of noise. The ideal weight parameter can be found by comparing the marginal likelihood of the model to the data distribution as shown in section 4.2.1. The Gaussian data used for training in all panels has a variance of five. The distributions learned by MCPC in (**a**) and (**b**) are obtained using MCPC’s spontaneous activity over 10,000 timesteps after training while maintaining the level of noise used during training. Moreover, the learning limit shown in (**b**) and (**c**) indicates where the variance of the Gaussian input layer of the MCPC model *σ*^2^*I* becomes larger than the variance of the data distribution.

## 3 Discussion

This work establishes how the brain could learn probability distributions of sensory inputs by relying solely on local computations and plasticity. We propose Monte Carlo predictive coding which is the first neural model that learns accurate probability distributions of sensory inputs using a hierarchical neural network with local computation and plasticity. MCPC introduces neural sampling to predictive coding using Langevin dynamics which enables: (i) the inference of full posteriors, (ii) the sampling of learned sensory inputs analogous to the brain imagining sensory stimuli, (iii) learning accurate generative models of sensory inputs, (iv) an ability to explain the variability of cortical activity and (v) learning robustly across noise types and intensities.

### 3.1 Benefits from computational abilities of MCPC

The identified neural dynamics of MCPC infer posterior distributions and generate data samples, and these abilities would provide great benefits to organisms supporting them. On one hand, the ability of MCPC to infer posterior distributions reflects the brain’s ability to infer statistically optimal representations of the environment. Such representations are key for survival through optimal perception [67] and decision-making [68]. On the other hand, our model’s ability to generate samples from learned sensory inputs is essential for offline replay. Cognitive functions that rely on offline replay include memory consolidation [69], planning of future actions [70], visual understanding [71], predictions [72], and decision-making [73]. Taken together, the neural activity of MCPC provides the basis upon which a wide array of other brain functions depend. This implies that MCPC might be useful not only for understanding generative learning, but also for unraveling the brain functions that potentially depend on its neural activity patterns.

### 3.2 Unified theory of cortical computation

MCPC integrates the strengths of predictive coding and neural sampling providing a unified theory of cortical computation.

MCPC, as a form of predictive coding, utilizes prediction error minimization for inference and learning in a hierarchical model. This alignment with predictive coding enables the application of PC’s potential cortical microcircuit implementations [74] and its implementation using dendritic errors [75] to MCPC. Additionally, MCPC can be applied to various learning tasks, similar to PC. For example, PC shows promising results in various classical tasks such as supervised learning, associative learning, representational learning, and reinforcement learning [27, 44, 76]. We expect MCPC to surpass PC in these tasks, owing to its enhanced inference dynamics that more closely approximate posterior distributions.

Concurrently, MCPC embodies neural sampling by employing neural dynamics to sample posterior distributions. Neural sampling was first proposed by Hoyer and Hyvärinen [10]. Since then, different implementations of sampling-based computations by the brain have been proposed [12, 29–34, 37]. These models have offered valuable insights that could be applied to MCPC. For instance, the sampling efficiency of MCPC could be improved through the use of excitatory and inhibitory recurrent networks, as suggested by Hennequin et al. [77]. Ultimately, MCPC opens new possibilities for a more comprehensive understanding of cortical computation and of the interplay between prediction-based learning and stochastic sampling mechanisms within the brain.

As a unified theory of cortical computation, MCPC can provide a unified account for a broad spectrum of cortical phenomena. This theory could bridge the explanatory scopes of both predictive coding and neural sampling. Predictive coding has played a pivotal role in providing a unified framework for explaining perception and attention [22]. It simultaneously offers insights into a range of neurological disorders such as schizophrenia, epilepsy, post-traumatic stress disorder, and chronic pain [23]. Predictive coding has also explained diverse neural phenomena ranging from retinal information encoding [24], alpha oscillations [25], and non-classical receptive fields [19]. Despite these results, PC has encountered challenges in explaining dynamic features of cortical activity. Neural sampling has provided significant insights in these areas. These features include the stimulus-dependence of neural variability [61, 78] and oscillations in the gamma band [79], strong transients at stimulus onset [80], and the spatiotemporal dynamics of bi-stable perception [6, 7]. By integrating PC with neural sampling, MCPC is poised to offer a comprehensive model capable of bridging the explanatory scopes of both predictive coding and neural sampling.

### 3.3 Relationship to other models introducing Langevin dynamics to brain-inspired generative models

Several brain-inspired generative models using Langevin dynamics have been proposed, and here we discuss their similarities and differences from MCPC.

Langevin dynamics were initially proposed as a mechanism through which the brain could infer posterior distributions via local neural computations, paving the way for models that elucidate diverse facets of perception and cortical functions [12, 29, 33, 37, 77]. Nevertheless, these models are either devoid of learning capabilities or rely on non-local plasticity mechanisms for weight adjustment. Additionally, the two experimental observations of neural variability captured by MCPC, as demonstrated in our work in figure 5, have not been previously modeled using Langevin dynamics. However, neural variability quenching at stimulus onset has been explained by other models of neural sampling [78]. Similarly, the increase in similarity between spontaneous and evoked neural activity has been explained by probabilistic generative learning in the brain [35]. Moreover, research has demonstrated that other characteristics of neural variability such as the relationship between neural variability and stimulus contrast can be effectively modeled through Langevin dynamics [37].

Langevin dynamics have also been applied in sparse coding models for posterior inference [31]. These models leverage local Langevin dynamics for inference and employ local plasticity rules specific to sparse coding for learning. When trained on patches of natural images, these models successfully learn simplecell receptive fields. Unlike MCPC, these sparse coding models do not possess hierarchical structures. Additionally, their learning capabilities have only been evaluated on relatively simple datasets, such as oriented bars.

Several machine learning studies have shown that generative models with Langevin dynamics learn accurate generative models of complex machine learning tasks [81, 82]. The studies show that the model with Langevin dynamics can outperform Variational Autoencoder and Generative adversarial networks on datasets such as MNIST, CIFAR-10, and CelebA. These studies confirm that models with Langevin dynamics can learn accurate generative models. However, in contrast to MCPC, these studies considered models that learn using non-local plasticity.

Since our initial presentation of MCPC [38], subsequent research [39, 40] has further validated that generative models employing Langevin dynamics learn precise generative models on complex tasks. Zahid et al. [39] also proposed the use of Langevin dynamics in predictive coding. However, their investigation focused on biologically implausible models with a singular latent layer, trained via backpropagation, diverging from MCPC’s approach. On the other hand, Dong and Wu [40] incorporated Langevin dynamics into generative models that leverage local computation and plasticity, showcasing capabilities for posterior inference and data generation using local neural dynamics akin to MCPC. Dong and Wu [40] employ exponential-family energy-based models, which differ from the hierarchical Gaussian models used in predictive coding. As a result, their proposed models are less directly linked to predictive coding than MCPC.

### 3.4 Experimental prediction

The core prediction of MCPC posits that the brain concurrently performs predictive error computations and sampling processes. Experimentation to substantiate MCPC would therefore involve detecting simultaneous prediction errors and neural sampling. According to predictive coding theories, prediction errors can be measured in the activity of error neurons, as discussed in this paper, or in the activity of dendrites [75]. Notably, this activity intensifies in response to unanticipated sensory inputs. Additionally, a measurable signature of sampling is a change in neural variability of value-encoding neurons as a result of a change in uncertainty associated with sensory inputs. An experimental approach to test MCPC’s prediction could, therefore, involve training animals to classify visual stimuli. Following their training, the experiment would measure the neural responses in the animals’ primary visual cortex when they are shown ambiguous and unambiguous stimuli. MCPC predicts that: (i) Neurons or dendrites that encode prediction errors will exhibit greater activity in response to ambiguous stimuli compared to non-ambiguous stimuli, and (ii) value-encoding neurons involved in sampling will display increased variability when processing ambiguous stimuli as opposed to unambiguous stimuli. An observed increase in both error-encoding activity and variability in value neurons in response to ambiguous stimuli, compared to unambiguous ones, would suggest the brain’s use of principles similar to those in MCPC for generative learning.

### 3.5 Limitations and Future work

#### 3.5.1 Extending MCPC to time-varying inputs

In this study, we focused on static sensory stimuli; however, the brain regularly processes temporally varying stimuli. Therefore, augmenting the MCPC model to predict future sensory inputs, would reflect brain function more accurately. By integrating elements from the temporal predictive coding model for sequential memory [83], we could develop a version of MCPC for learning generative models of time-varying sensory inputs. This adaptation is expected to further align MCPC with the dynamic processing capabilities of the brain without compromising its biological plausibility.

#### 3.5.2 Improving the sampling speed of Langevin dynamics

Despite the promising results in this study, sampling using MCPC’s Langevin dynamics requires long inference times [45]. The inference duration could be significantly shortened by relying on advancements in machine learning. For instance, adding higher-order terms to the Langevin dynamics, such as momentum, dramatically improves convergence speed [84]. Ma et al. [85] also proposed a general framework for improving the sampling efficiency of Langevin-based sampling. By applying these principles to MCPC, the sampling speed is expected to increase, reaching a value that resembles the fast sampling of the brain [1]. Importantly, the learning performance of MCPC is then anticipated to improve, as the inferences will capture the posterior more effectively.

#### 3.5.3 Mapping MCPC’s noise to sources of noise in the brain

Currently, mapping the noise variable in MCPC’s neural dynamics to distinct noise sources in the brain remains a challenge. Cortical circuits have various forms of stochasticity that could support the random dynamics of MCPC [86]. However, the constraints on the noise variable within the dynamics of MCPC are notably minimal. MCPC does not mandate that the noise follow a particular distribution, nor does it specify a required noise level. Consequently, predicting which types of noise in the brain could facilitate the stochastic dynamics of MCPC proves challenging. To establish a more direct link between MCPC’s noise and its potential physiological origins, a spiking implementation of MCPC could be identified. Rethinking MCPC using spiking neural networks, as done in the spiking models of predictive coding [87], might add constraints on the location and type of noise needed. These constraints could create a clear connection to the physiological sources of noise.

## 4 Methods

### 4.1 Models

In this paper, we compare three methods for learning hierarchical Gaussian models: Monte Carlo predictive coding, predictive coding, and backpropagation in a deep latent Gaussian model.

#### 4.1.1 Monte Carlo predictive coding

Algorithm 1 shows the complete implementation of MCPC used for all the simulations in the paper. This algorithm is a discrete-time equivalent of MCPC where the dynamics of MCPC given in equation (3) are discretized using the Euler–Maruyama method. Moreover, the algorithm contains a MAP inference before the MCPC inference to shorten MCPC’s mixing time during inference.

#### 4.1.2 Predictive coding

We briefly review the predictive coding framework and its implementation used in this paper. Following the formulation of predictive coding by Bogacz [21] which we refer to with PC, predictive coding learns a hierarchical Gaussian model. The model is learned by iterating over two steps that minimise the joint log-likelihood 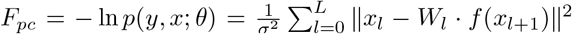 where *x*_0_ is clamped to an observation *y*. First, PC uses neural dynamics that follow the gradient flow on *F*_*pc*_ to infer the Maximum a-posteriori estimate of the latent states conditioned on the observation:

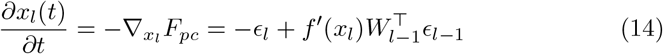

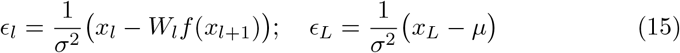

##### Algorithm 1: Monte Carlo predictive coding (MCPC)

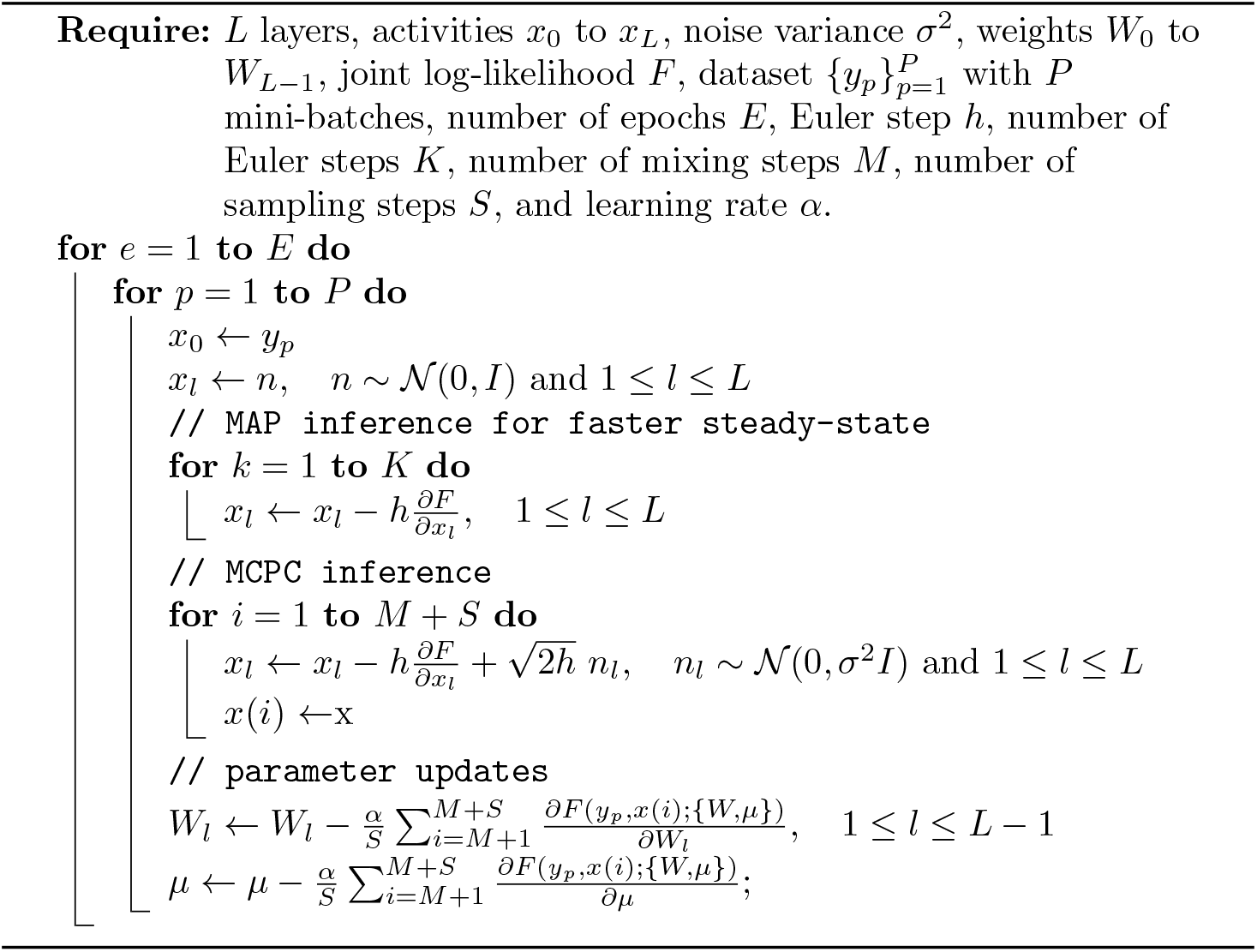

##### Algorithm 2: Predictive coding (PC)

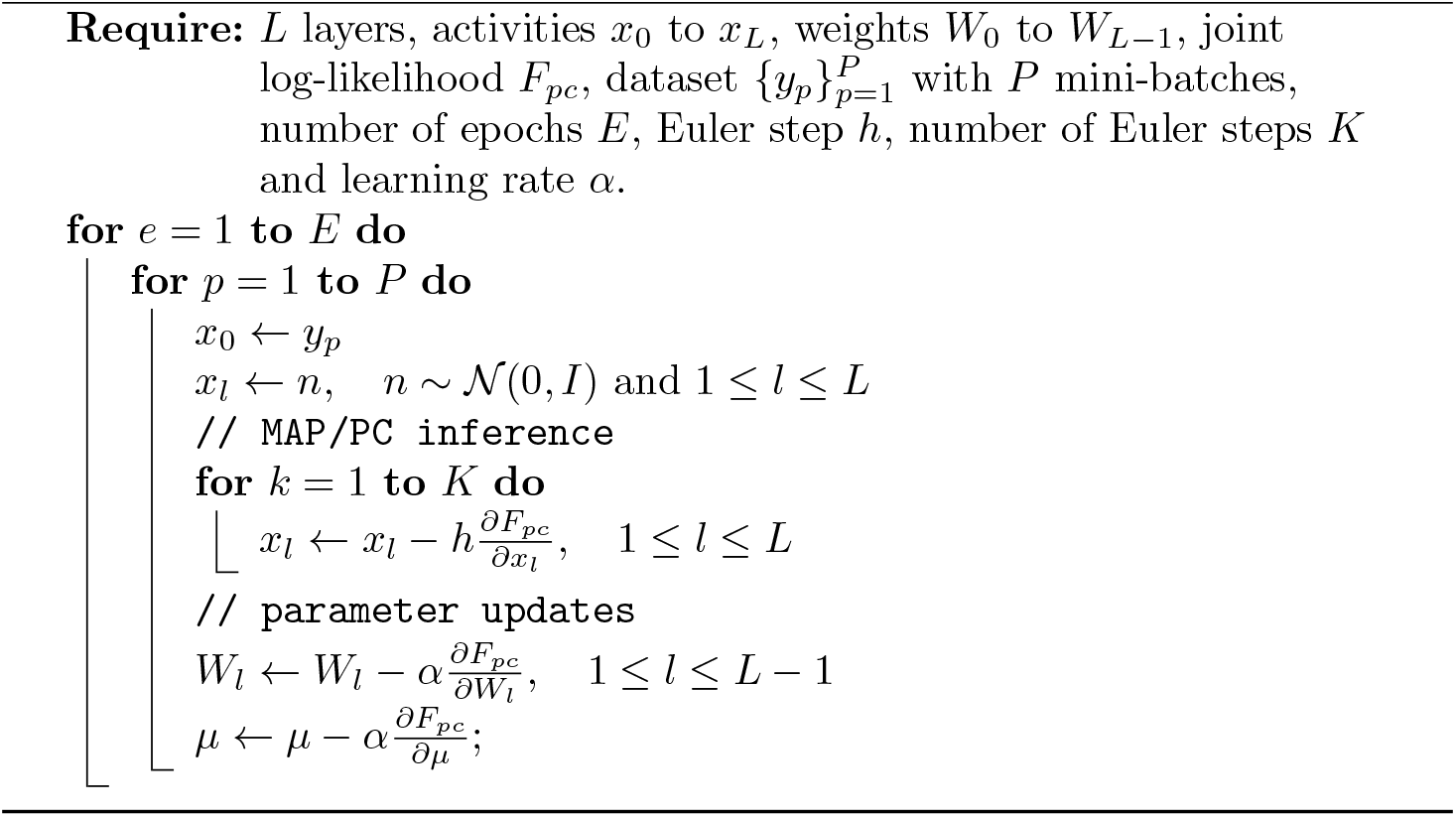

Second, PC updates the parameters with a gradient step on *F*_*pc*_, evaluated on the converged MAP estimate *x*\* with error *ϵ*\*:

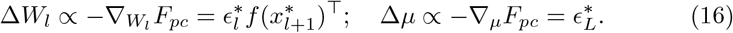

These computations can be implemented in the same neural network with local computation and plasticity as MCPC. This is because PC only differs from MCPC through an additional noise term in the neural dynamics and an integral in the weight updates. Algorithm 2 shows the complete implementation of PC used for all the simulations in the paper. This algorithm is a discrete-time equivalent of PC where the dynamics of PC given in equation (14) are discretized using Euler’s method which consists of taking small discrete steps in the derivative direction.

#### 4.1.3 Deep latent Gaussian models

We implement DLGMs as a benchmark model because they are the standard machine learning model for learning hierarchical Gaussian models. DLGMs were first proposed by Rezende et al. [51] and they consist of two main components: a generative model and an inference model. Each of these models is represented by separate neural networks. The generative model is responsible for generating samples, while the inference model approximates the posterior distribution over the latent variables given the observed data. To train this model, we utilize the reparameterization trick [41, 51]. This technique allows for the backpropagation of gradients through stochastic nodes, enabling efficient and accurate gradient-based optimization to learn the parameters of both the generative and inference models. However, DLGMs are not a biologically plausible model of generative learning in the brain. One major shortcoming is that the plasticity mechanisms used by DLGMs are not local. This study uses a modified version of the DLGMs implementation by Zhuo [88]. The implementation is modified so that the inference network learns a rank 1 approximation of the covariance matrices of the posterior. This reduces the number of parameters of the inference network without significantly affecting the learning performance of DLGMs. To ensure a fair comparison with PC and MCPC, we employ DLGMs with generative models possessing a parameter count equivalent to that of PC and MCPC. Furthermore, the inference networks of DLGMs are constrained to maintain a parameter count equal to their generative counterparts.

### 4.2 Learning tasks

Throughout the paper two generative learning tasks are studied: a Gaussian learning task and a handwritten digit image learning task.

#### 4.2.1 Gaussian learning task

In this task, the data has a Gaussian distribution that can be learned by the model in figure 1a. This model is tractable, facilitating a comparison of MCPC’s steady state inference with the marginal likelihood *p*(*y*; *θ*) and the posterior distribution *p*(*x*| *y*; *θ*). The model’s tractability also enables a direct comparison of the optimal parameters to the parameters learned by MCPC and PC. Equations (17) and (18) provide the data distribution and the model used for this task.

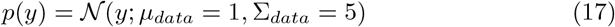

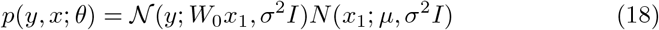

The marginal likelihood and the posterior distributions are given in equations (19) and (20).

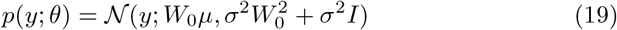

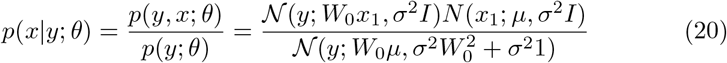

To evaluate steady-state neural activity with and without inputs, we employ 10,000 inference steps for MCPC and 2,000 steps for PC. The optimal parameter values, 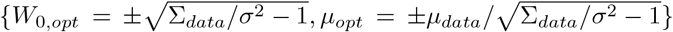, are identified by comparing the marginal likelihood to the data distribution. We train an MCPC and a PC model on this task using the parameters in table 2.

**Table 2.** Parameters of MCPC, PC, and DLGMs. The parameters are given for the linear task and the hyperparameter search space for the MNIST task. The parameters under MCPC Inference are only used for MCPC inference. The parameters under PC Inference are used for PC inference and the MAP inference in algorithm 1 implementing MCPC. The parameters under Architecture and Learning are shared by MCPC, PC, and DLGMs.

|  | Gaussian | MNIST |
| --- | --- | --- |
| MCPC Inference |  |  |
| **optimizer** | SGD | SGD |
| **lr** | 0.02 | {0.003, 0.01, 0.03, 0.1, 0.3} |
| mixing steps $T$ | 150 | 50 |
| sampling steps $M$ | 1 | 100 |
| PC Inference |  |  |
| **optimizer** | Adam | Adam |
| **lr** | 0.02 | {0.03, 0.1, 0.3, 0.7} |
| **max_steps** | 150 | 250 |
| Architecture |  |  |
| input dimension | 1 | 748 |
| number of latent layers $L$ | 1 | 3 |
| dimension of $x_1$ to $x_{L-1}$ | - | {128, 256, 360} |
| dimension of $x_L$ | 1 | {10, 15, 20, 25, 30} |
| activation function | linear | {ReLU, tanh} |
| noise variance $\sigma^2$ | 1 | 1 |
| Learning |  |  |
| **optimiser** | Adam | Adam |
| **lr** | 0.02 | {0.001, 0.003, 0.01, 0.03} |
| **decay** | 0 | {0, 0.01, 0.1, 1} |
| **num_epochs** | 75 | 50 |
| **batch_size** | 256 | {64, 128, 256} |

#### 4.2.2 MNIST learning task

In this task, the dataset comprises 28×28 binary images representing handwritten digits across ten categories. The model architecture and training parameters used for this task are determined using a hyperparameter search. Moreover, the model has been adapted to have a Bernoulli input layer. In contrast to the Gaussian learning task, the model is intractable due to its hierarchical structure and non-linear activation functions, necessary for accurate learning. Consequently, direct assessment of inferences and model parameters is not feasible. Instead, we visualize the neural activity of MCPC and PC with and without inputs, we compare the neural activity of MCPC and PC with inputs to the inferences of an artificial ideal observer, and we measure the learning performance of the models using three metrics.

##### Model parameters

The model architecture and training parameters used for this task are determined using a hyperparameter search summarised in table 2. The dataset includes 60,000 training images and 10,000 testing images, of which 6,000 images are used for hyperparameter tuning and 4,000 for evaluation. Supplementary tables 1, 2, and 3 compile the search results.

##### Bernoulli input layer

The model used for this task has been adapted to have a Bernoulli input layer. For binary images, the model’s input layer is transformed into a multivariate Bernoulli distribution. This modification yields the joint log-likelihood 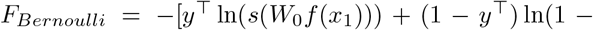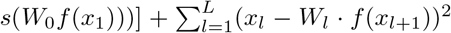 where *s* is a sigmoid function. This change does not compromise the biological plausibility of MCPC because it only introduces an additional non-linearity in the inference dynamics of *x*_1_ and the parameter update for *W*_0_. This modification is shown below:

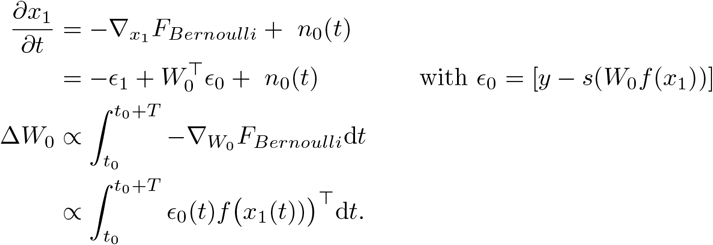

#### Visualisation of MCPC’s and PC’s neural activity

We visualize MCPC’s and PC’s neural activity to assess the inference with and without inputs. We record and display the input neurons’ activity over time to visualize inferences without inputs. To visualize with inputs, we use a linear classifier that decodes the digit class distribution from the latent state *x*_*L*_. The classifier is trained on full images of the training data to transform MAP inferences of the latent state *x*_*L*_ to the corresponding digit classes. The digit classes are then assigned coordinates using a convex combination of 10 evenly spaced points on a unit circle [49], resulting in a two-dimensional visualization. In figure 2, we visualize the inference for a full image part of the evaluation data and a partially masked version of the same image. For both visualizations, we use 10,000 inference steps for MCPC and 2,000 for PC.

#### Quantification of neural activity of MCPC and PC with inputs

We compare the neural activity inferred by MCPC and PC with inputs to the posterior inferred by an artificial ideal observer. This comparison quantifies how well MCPC and PC approximate the posterior distribution. The artificial ideal observer is a ResNet-9 classifier. We employ the same linear classifier as used for the visualizations to decode a digit class distribution from the neural activity of MCPC and PC. The decoded class distribution can then be compared to the digit class distribution inferred by the ideal observed. For PC, the digit class distribution is obtained by decoding the inferred latent state *x*_*L*_ at convergence. For MCPC, this distribution is obtained by decoding the fluctuating latent state *x*_*L*_ at steady state and averaging the decoded distributions across MCPC samples. MCPC and PC are compared to the ideal observer by computing the KL divergence between the digit class distributions on the MNIST evaluation set with the top half of the images masked. For the random baseline, we compute the Kullback-Leibler divergence between the posterior distribution inferred by the ideal observer and the distributions inferred by both MCPC and PC, after these have been randomly shuffled. This shuffling results in the distributions inferred by PC and MCPC being associated with random inputs.

#### Performance metrics

We assess the learning accuracy of MCPC and PC using three metrics. Firstly, the Frechet Inception Distance (FID) evaluates the quality and diversity of generated images [56]. The FID is computed by comparing the evaluation images with 5000 generated images using a public FID implementation [89]. Secondly, we approximate the marginal log-likelihood for the evaluation images to assess a model’s generalization performance. The marginal log-likelihood is approximated using the following Monte Carlo estimate from 5000 latent state samples:

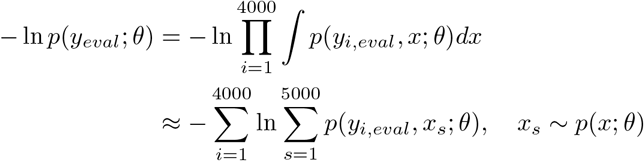

The samples to compute both the FID and the marginal log-likelihood are obtained using ancestral sampling. Finally, the reconstruction MSE measures the error between images and the images reconstructed by a model when inputted with the bottom half of the images. We calculate the error as the mean squared error between the top half of the original and reconstructed images for the MNIST evaluation set. The reconstructed images are obtained by performing MAP inferences of the missing image pixel values.

### 4.3 Measuring cortical-like properties of MCPC’s neural activity

We measure two properties of MCPC models: the neural variability at stimulus onset, and the similarity between spontaneous and average evoked activity during training.

#### 4.3.1 Evaluating neural variability

We first measure the temporal neural variability of the latent state activity at stimulus onset for MCPC and PC. This mimics the neural variability recording in the V1 region of cats done by Churchland et al. [61]. The model used for this experiment is trained on MNIST with MCPC to optimize the model’s FID. Moreover, we measure the neural variability around 256 stimuli onsets from the MNIST evaluation set. We measure the temporal neural variability by computing the standard deviation of the activity of the latent states over a sliding window of 1000 timesteps. Then, we average the neural variability over all latent states for the 256 stimuli onsets. Churchland et al. [61] employed a 50-ms sliding window to estimate the variance in membrane potential of individual neurons. Then, they averaged the neural variability across the 52 recorded neurons and all stimuli to plot the mean change in neural variability at stimulus onset. Our approach replicates the experimental approach of Churchland et al. [61] for measuring neural variability of membrane potentials from cat V1.

#### 4.3.2 Similarity of spontaneous and average evoked neural activity

Our method to measure the similarity of spontaneous and average evoked activity follows the approach used to measure this similarity in the V1 region of ferrets. Berkes et al. [35] recorded the spontaneous and average evoked neural activity with a linear array of 16 electrodes implanted in V1 of 16 ferrets (approximately 4 ferrets per age group). They measured the evoked activity for natural stimuli, sinusoidal gratings, and random noise. Moreover, they quantified the similarity in activity using the KL divergence. Our approach relies on similar sensory inputs and uses the same quantification metric. We measure the similarity between the spontaneous activity and the average evoked activity using a KL divergence in 10 MCPC models. We compute the KL divergence at different steps during training on MNIST as follows: First, we record the evoked activity of an MCPC model to (i) 256 samples from MNIST’s evaluation set (natural stimuli), (ii) 256 samples of sinusoidal gratings with 16 possible orientations, and (iii) 256 samples of random binary images. We record from five randomly selected latent states in the first latent layer (*l* = 1) for 9,500 inference steps. Recording from five latent states reduces the computation time while maintaining representative results for the whole network. Moreover, recording from the first latent layer mirrors the V1 region which is the first cortical area that processes visual information. Second, we record the spontaneous activity of the model for the same five latent states for 9,500 activity updates. Third, for each type of evoked activity, we compute the average experimental distribution of evoked activities across samples. Finally, we compare the three average distributions to the distribution of spontaneous activity using the KL divergence as implemented by Pérez-Cruz [90]. We repeat this procedure for 10 models trained on MNIST with MCPC to optimize the model’s FID. These models have the same architecture and learning parameters but they are initialized using a different seed. After, the KL divergence for natural stimuli is compared using paired samples t-tests to the KL divergence for gratings and for noise.

## 5 Data availability

The MNIST dataset [42] is publicly available.

## 6 Code availability

The codebase with all models and experiments can be found at the following link: https://github.com/gaspardol/MonteCarloPredictiveCoding.git. This code is based on the implementation of predictive coding by Song et al. [44].

## 7 Acknowledgement

This work has been supported by the Medical Research Council UK grant MC UU 00003/1.

