## Supplementary information for "Learning probability distributions of sensory inputs with Monte Carlo Predictive Coding"

#### 1 Encoding the variance of Gaussian model layers in MCPC

In the hierarchical Gaussian model, layer variance $\sigma^2$ can be represented using two distinct methods within MCPC. First, it may be integrated into the joint log-likelihood $F$, as is common in predictive coding approaches [1, 2]. Alternatively, it can be included in the noise variance of the MCPC inference dynamics. Both methods are possible because the steady-state distribution of MCPC's neural dynamics depends on the ratio between the energy function being minimized and the noise variance, as illustrated in proposition 1 of the main text. Consequently, the layer variance can either be removed from the energy function $F$ and incorporated into the noise variance or retained in the energy function with the noise variance set to a standard value. For this research, we opted for the former approach. This selection enhances clarity by explicitly presenting the connection between noise variance and model variance. It also highlights an additional mechanism in MCPC for representing the variance of its generative model, which is not present in predictive coding.

#### 2 Learning in a linear model with one input neuron and one latent state using MCPC and PC

We compare training the linear model given in figure 1a of the main text using MCPC and PC. We find that MCPC only has equilibrium points at the optimal model parameters while PC does not have an equilibrium for finite weights. This demonstrates the theoretical superiority of MCPC over PC. Additionally, this elucidates why the weights in PC become excessively large which is observed across tasks. Here, we consider a linear model with identity covariance matrices for simplicity. However, the result generally holds for any covariance matrices.

##### 2.1 Learning with MCPC

**Inference.** MCPC infers the posterior distribution $p(x_1|y;\theta)$ which for the considered model equals:

$$p(x_1|y;\theta) = \frac{p(y,x;\theta)}{p(y;\theta)} = \frac{\mathcal{N}(y;Wx_1,I)N(x_1;\mu,I)}{\mathcal{N}(y;W\mu,W^2+1)}$$
$$= \mathcal{N}\Big(x_1;\frac{W_0y+\mu}{W_0^2+1},\frac{1}{W_0^2+1}\Big).$$

**Expected parameter updates.** The expected parameter updates can be found using MCPC's equation for parameter updates given in equations (6) and

(7) of the main text. For the weight $W_0$, the expected parameter update is given by:

$$\begin{aligned}\mathbb{E}\{\Delta W_0\} &\propto \mathbb{E}\{\mathbb{E}_{p(x_1|y;\theta)}\{e_0^* x_1^*\}\} = \mathbb{E}\Big\{\int (y - W_0 x) p(x_1|y;\theta) dx_1\Big\} \\ &= \mathbb{E}\Big\{\int (y - W_0 x)\mathcal{N}\Big(x_1; \frac{W_0 y + \mu}{W_0^2 + 1}, \frac{1}{W_0^2+1}\Big) dx_1\Big\} \\ &= \frac{1}{(w_0^2+1)^2}\Big( - W_0(Var\{y\} + \mathbb{E}\{y\}^2) + \mu(W_0^2 - 1)\mathbb{E}\{y\} + W_0^3 + W_0\mu^2 + W_0\Big).\end{aligned}$$

For the prior mean $\mu$, the expected parameter update is given by:

$$\begin{aligned}\mathbb{E}\{\Delta\mu\} &\propto \mathbb{E}\{\mathbb{E}_{p(x_1|y;\theta)}\{\epsilon_1^*\}\} = \mathbb{E}\Big\{\int (x_1^* - \mu) p(x_1|y;\theta) dx_1\Big\} \\ &= \mathbb{E}\Big\{\int (x_1^* - \mu)\mathcal{N}\Big(x_1; \frac{W_0 y + \mu}{W_0^2+1}, \frac{1}{W_0^2+1}\Big) dx_1\Big\} \\ &= -\frac{W_0}{W_0^2+1}\Big(\mathbb{E}\{y\} - W_0\mu\Big).\end{aligned}$$

In these expressions, $\mathbb{E}\{y\}$ equals the mean of the data and $Var\{y\}$ equals the variance of the data.

**Nullclines.** The nullclines of the model can be found by equating the expected parameter updates to zero. Therefore, the nullclines for $\mathbb{E}\{\Delta W_0\} = 0$ are given by the expression:

$$\mu = \frac{-(W_0^2 - 1)\mathbb{E}\{y\} \pm \sqrt{(W_0^2-1)^2\mathbb{E}\{y\}^2 + 4W_0^2(Var\{y\} + \mathbb{E}\{y\}^2 - W_0^2 - 1)}}{2W_0}.$$

Similarly, the nullcline for $\mathbb{E}\{\Delta\mu\} = 0$ is given by the expression:

$$\mu = \frac{\mathbb{E}\{y\}}{W_0}.$$

**Equilibrium points.** Equilibrium points lie at the intersection between the nullclines $E\{\Delta W\} = 0$ and the nullcline $E\{\Delta\mu\} = 0$. By equating the nullclines, you obtain the condition on equilibrium points:

$$W_0^2 = Var\{y\} + 1.$$

The parameter values of the equilibrium points are therefore $\{W_0 = \pm\sqrt{Var\{y\} - 1}, \mu = \pm\frac{\mathbb{E}\{y\}}{\sqrt{Var\{y\}-1}}\}$. These values are optimal because a model with these parameter values has a marginal likelihood $p(y;\theta)$ equal to $\mathcal{N}(y; \mathbb{E}\{y\}, Var\{y\})$.

#### 2.2 Learning with PC

**Inference.** For the linear model, PC inference using the dynamics given in equation (14) of the main text converges to the latent state $x_1^*$ for a given input

$y$.

$$\frac{\partial x_1^*(t)}{\partial t} = 0 = -\epsilon_1^* + W_0\epsilon_0^* = -(x_1^* - \mu) + W_0(x_0^* - W_0 x_1^*)$$

$$x_1^* = \frac{\mu + W_0 y}{1 + W_0^2}$$

**Expected parameter updates.** The expected parameter updates can be found using PC's equation for parameter updates given in equation (16) of the main text. For the weight $W_0$, the expected parameter update is given by:

$$\begin{aligned}
\mathbb{E}\{\Delta W_0\} &\propto \mathbb{E}\{e_0^* x_1^*\} = E\{(y - W_0 x_1^*)x_1^*\} \\
&= \mathbb{E}\Big\{\Big(y - W_0\frac{\mu + W_0 y}{1 + W_0^2}\Big)\frac{\mu + W_0 y}{1 + W_0^2}\Big\} \\
&= \frac{1}{(1 + W_0^2)^2}\Big(W_0\mathbb{E}\{y^2\} + (\mu - W_0^2\mu)\mathbb{E}\{y\} - W_0\mu^2\Big) \\
&= \frac{1}{(1 + W_0^2)^2}\Big(W_0(Var\{y\} + \mathbb{E}\{y\}^2) + \mu(1 - W_0^2)\mathbb{E}\{y\} - W_0\mu^2\Big).
\end{aligned}$$

For the prior mean $\mu$, the expected parameter update is given by:

$$\begin{aligned}
\mathbb{E}\{\Delta\mu\} \propto \mathbb{E}\{\epsilon_1^*\} = \mathbb{E}\{x_1^* - \mu\} &= \mathbb{E}\{\frac{\mu + W_0 y}{1 + W_0^2} - \mu\} \\
&= \frac{\mu + W_0}{1 + W_0^2}\mathbb{E}\{y\} - \mu.
\end{aligned}$$

**Nullclines.** The nullclines for $\mathbb{E}\{\Delta W_0\} = 0$ are given, therefore, by the expression:

$$\mu = \frac{-(W_0^2 - 1)\mathbb{E}\{y\} \pm \sqrt{(W_0^2 - 1)^2\mathbb{E}\{y\}^2 + 4W_0^2(Var\{y\} + \mathbb{E}\{y\}^2)}}{2W_0}.$$

Similarly, the nullcline for $\mathbb{E}\{\Delta\mu\} = 0$ is given by the expression:

$$\mu = \frac{\mathbb{E}\{y\}}{W_0}.$$

**Equilibrium points.** We can attempt to find equilibrium points by finding the intersection between the nullclines for $\mathbb{E}\{\Delta W_0\} = 0$ and the nullcline for$\mathbb{E}\{\Delta\mu\} = 0$ as follows:

$$\begin{aligned}
&\frac{\mathbb{E}\{y\}}{W_0} = \frac{-(1 - W_0^2)\mathbb{E}\{y\} \pm \sqrt{(1 - W_0^2)^2\mathbb{E}\{y\}^2 + 4W_0^2(Var\{y\} + \mathbb{E}\{y\}^2)}}{-2W_0} \\
&\mathbb{E}\{y\}\Big(-1 - W_0^2\Big) = \pm\sqrt{\mathbb{E}\{y\}^2(1 - W_0^2)^2 + 4W_0^2(Var\{y\} + \mathbb{E}\{y\}^2)} \\
&\mathbb{E}\{y\}^2\Big(1 + W_0^4 + 2W_0^2\Big) = \mathbb{E}\{y\}^2(1 + W_0^4 - 2W_0^2) + 4W_0^2(Var\{y\} + \mathbb{E}\{y\}^2) \\
&4W_0^2\mathbb{E}\{y\}^2 = 4W_0^2(Var\{y\} + \mathbb{E}\{y\}^2) \\
&Var\{y\} = 0
\end{aligned}$$

There are therefore no equilibrium points except when $Var\{y\} = 0$ where all the points on $\mu = \frac{\mathbb{E}\{y\}}{W}$ are fixed points. Consequently, the parameter updates of PC do not converge.

### 3 Predictive coding optimizes an infinitely loose bound on the marginal log-likelihood $\ln p(y;\theta)$

Our results suggest that predictive coding fails to learn accurate generative models. This result could be explained by a theoretical shortcoming of predictive coding which is also present in a range of other theories of learning in the brain as discussed below.

Predictive coding learns a generative model by changing its model parameters to maximize the marginal likelihood:

$$p(y;\theta) = \int p(y,x;\theta)\mathrm{d}x$$

Predictive coding uses the variational expectation-maximization algorithm to optimize this marginal which is intractable [1, 3–5]. This algorithm employs the variational distribution $q(x;\phi)$, parametrized by $\phi$, to estimate the posterior $p(x|y;\theta)$. It then establishes the free energy $\mathcal{F}$ as an upper bound on the negative log-likelihood:

$$\begin{aligned}\mathcal{F}(\phi,\theta) &= -\mathbb{E}_q\{\ln p(y,x;\theta)\} + \mathbb{E}_q\{\ln q(x;\phi)\} \\ &= -\ln p(y;\theta) + D_{\mathrm{KL}}(q(x;\phi)\|p(x|y;\theta)) \geq -\ln p(y;\theta),\end{aligned} \tag{1}$$

with $D_{\mathrm{KL}}$ the KL divergence which is always positive.

The variational expectation-maximization algorithm learns by iterating over two steps. First, it minimizes the free energy $\mathcal{F}$ w.r.t. $\phi$ for the current parameters $\theta$. In other words, it finds an approximation $q(x;\phi)$ for the posterior. Second, the algorithm minimizes the free energy w.r.t. $\theta$ for the parameters $\phi$ inferred in the first step.

Predictive coding uses the Dirac delta distribution $q(x;\phi) = \delta(x-\phi)$ as variational distribution [1], enabling an implementation of predictive coding using local computation and plasticity. Importantly, the entropy $-\mathbb{E}_q\{\ln q(x;\phi)\}$ for this variational distribution is equal to minus infinity. The free energy $\mathcal{F}$ minimized by predictive coding is therefore infinite, making it an infinitely loose bound on the marginal likelihood.

This infinitely loose bound raises doubts about the significance of the objective minimized by predictive coding. This theoretical shortcoming may explain the observed poor learning performance of predictive coding in this study. Moreover, a range of theories for learning in the brain [6–8] are based on a similar energy function as PC's variational free energy, importing the problem of implicitly ignoring an infinite entropy. Future work could therefore explore whether these other learning algorithms exhibit similar suboptimal learning capabilities to those observed in predictive coding.

### 4 Proofs for MCPC capturing the variability of cortical activity

#### 4.1 Neural variability decreases at stimulus onset

At stimulus onset, the variance of MCPC's steady-state activity decreases. This is because, MCPC's neural activity of the latent state switches from representing the marginal distribution $p(x;\theta)$ to the posterior $p(x|y;\theta)$, which in expectation has a lower variance. The steady state of MCPC's latent state before stimulus onset can be derived similarly to proposition 2 of the main text and equals the marginal distribution $p(x;\theta)$:

$$p^{ss}(x) = \int \frac{e^{-F/\sigma^2}}{Z} dx = \int \frac{e^{\ln p(y=x_0,x;\theta)}}{Z} dx_0 = p(x;\theta) \int \frac{p(x_0|x;\theta)}{Z} dx_0$$
$$= p(x;\theta)$$

Moreover, the law of total variance presented in equation (2) shows that the posterior $p(x|y;\theta)$ has in expectation a lower variance than the marginal distribution $p(x;\theta)$. Therefore, the neural variability of MCPC's latent state is expected to decrease at stimulus onset as experimentally observed in the brain.

$$Var[p(x;\theta)] = E\{Var[p(x|y;\theta)]\} + Var[E\{p(x|y;\theta)\}] \tag{2}$$

#### 4.2 Natural stimuli specific increase in similarity between spontaneous and evoked neural activity

As the training progresses, the similarity between MCPC's spontaneous activity (encoding the marginal distribution $p(x;\theta)$) and its average evoked activity increases (encoding the posterior distribution $p(x|y;\theta)$). This increase in similarity, however, is specific to the natural stimuli used during training. It does not extend to data outside the training set's distribution, indicating a specialized adaptation to the learned stimuli. This phenomenon can be explained by Bayesian statistics. The marginal distribution $p(x;\theta)$ of a latent variable model with latent state $x$ and input $y$ serves as the model's prior distribution over the latent state, encapsulating prior expectations. Initially, the model parameters are randomly initialized, leading to a discrepancy between the model's average posterior for its inputs and its prior distributions. However, as training advances, the model refines its generative capabilities, aligning the marginal likelihood $p(y;\theta)$ closer to the data distribution $p(y)$. Consequently, the model's prior over the latent state begins to mirror the average posterior distribution more accurately. If the model achieves perfect learning, its prior will match exactly with the expected posterior for the training data. This result follows from Bayesian statistics that specifies that the probability distribution $p(x)$ equals $\int p(x|y)p(y)dy$ (see equation 3). However, the prior of the model is specific to the data distribution. Consequently, when considering posterior distributions for data samples outside the training data distribution, the prior will not equate to the average posterior.

$$p(x) = \int p(x,y)dy = \int p(x|y)p(y)dy \tag{3}$$

### 5 Hyperparameter search results

This section summarises the optimal model parameters for MCPC, PC, and DLGMs found during the hyperparameter search to maximise the FID, the marginal likelihood $-\ln p(y_{eval})$, and the reconstruction MSE.

| | MCPC | PC | DLGM |
|---|---|---|---|
| Architecture | | | |
| input dimension | 748 | 748 | 748 |
| number of hidden layers | 3 | 3 | 3 |
| hidden layers dimension | (128, 128, 20) | (128, 128, 20) | (256,256,20) |
| activation function | ReLU | ReLU | ReLU |
| PC Inference | | | |
| `optimizer` | Adam | Adam | - |
| `lr` | 0.7 | 0.1 | - |
| `max_steps` | 250 | 250 | - |
| MCPC Inference | | | |
| `optimizer` | SGD | - | - |
| `lr` | 0.1 | - | - |
| mixing steps | 50 | - | - |
| sampling steps | 100 | - | - |
| Learning | | | |
| `optimiser` | Adam | Adam | Adam |
| `lr` | 0.01 | 0.01 | 0.001 |
| `decay` | 0.0 | 0.1 | 0.01 |
| `num_epochs` | 50 | 50 | 50 |
| `batch_size` | 256 | 128 | 128 |

**Table 1.** Results of the hyperparameter search to maximize the FID for MCPC, PC, and DLGMs.

| | MCPC | PC | DLGM |
|---|---|---|---|
| Architecture | | | |
| input dimension | 748 | 748 | 748 |
| number of hidden layers | 3 | 3 | 3 |
| hidden layers dimension | (128, 128, 20) | (128, 128, 25) | (128, 128, 10) |
| activation function | ReLU | Tanh | ReLu |
| PC Inference | | | |
| `optimizer` | Adam | Adam | - |
| `lr` | 0.1 | 0.3 | - |
| `max_steps` | 250 | 250 | - |
| MCPC Inference | | | |
| `optimizer` | SGD | - | - |
| `lr` | 0.03 | - | - |
| mixing steps | 50 | - | - |
| sampling steps | 100 | - | - |
| Learning | | | |
| `optimiser` | Adam | Adam | Adam |
| `lr` | 0.003 | 0.001 | 0.001 |
| `decay` | 0.1 | 0.01 | 0.01 |
| `num_epochs` | 50 | 50 | 50 |
| `batch_size` | 64 | 256 | 256 |

**Table 2.** Results of the hyperparameter search to maximize the marginal likelihood for MCPC, PC, and DLGMs.

### 6 Supplementary videos

**Video 1. MCPC posterior inference in linear model.** Animation of the activity of the latent state in a linear model with one latent state during MCPC inference for a constant input. In this animation, the orange dot shows the time-varying activity of the latent state. The blue histogram summarises the activity of the latent state from the beginning of the animation to the time point in the animation being visualized. Finally, the black curve shows the true posterior distribution that can be analytically calculated from the model parameters and the input to the model.

**Video 2. MCPC posterior inference in non-linear model for half masked MNIST digit.** Animation of the activity of the latent layer $x_L$ in a non-linear model trained on the MNIST dataset during MCPC inference for a half-masked digit input. The orange dot shows the time-varying activity of the latent state $x_L$ transformed to coordinates using a linear classifier and a convex combination of 10 evenly spaced points on a unit circle. The linear classifier is trained to decode digit class distributions from the latent state $x_L$. The decoded class distribution can then be transformed to a coordinate using the convex combination. The blue hexagons show the probability density of a

| | MCPC | PC | DLGM |
|---|---|---|---|
| Architecture | | | |
| input dimension | 748 | 748 | 748 |
| number of hidden layers | 3 | 3 | 3 |
| hidden layers dimension | (256, 256, 20) | (256, 256, 30) | (256, 256, 20) |
| activation function | ReLU | Tanh | ReLu |
| PC Inference | | | |
| `optimizer` | Adam | Adam | - |
| `lr` | 0.7 | 0.7 | - |
| `max_steps` | 250 | 250 | - |
| MCPC Inference | | | |
| `optimizer` | SGD | - | - |
| `lr` | 0.03 | - | - |
| mixing steps | 50 | - | - |
| sampling steps | 100 | - | - |
| Learning | | | |
| `optimiser` | Adam | Adam | Adam |
| `lr` | 0.001 | 0.003 | 0.003 |
| `decay` | 0.01 | 0.1 | 0.1 |
| `num_epochs` | 50 | 50 | 50 |
| `batch_size` | 64 | 256 | 128 |

**Table 3.** Results of the hyperparameter search to maximize the reconstruction MSE for MCPC, PC, and DLGMs.

two-dimensional histogram of the activity of the latent state from the beginning of the animation to the time point in the animation being visualized.

**Video 3. MCPC posterior inference in non-linear model for full MNIST digit.** Animation of the activity of the latent layer $x_L$ in a non-linear model trained on the MNIST dataset during MCPC inference for a full digit input. The orange dot and the blue hexagons have been determined as described in supplementary video 2.

**Video 4. MCPC unclamped activity of sensory input neuron in a linear model.** Animation of the activity of the input neuron in a linear model with one latent state and one input neuron resulting from MCPC dynamics when the input neuron is unclamped. The orange dot shows the input neuron activity over time. The histogram summarises the activity of the input state from the beginning of the animation to the time point in the animation being visualized. The black curve shows the marginal likelihood that can be analytically calculated from the model parameters.

**Video 5. MCPC unclamped activity of sensory input neurons in non-linear model trained on MNIST.** Animation of the activity of the input neurons in a non-linear model trained on MNIST resulting from MCPC dynamics when the input neurons are unclamped.
