## Supplementary figures and images for "Learning probability distributions of sensory inputs with Monte Carlo Predictive Coding"

### Supplementary video 1

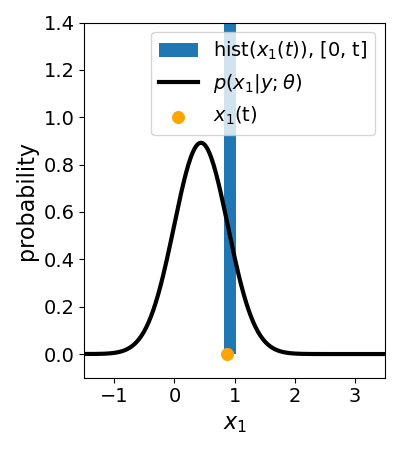

### Supplementary video 2

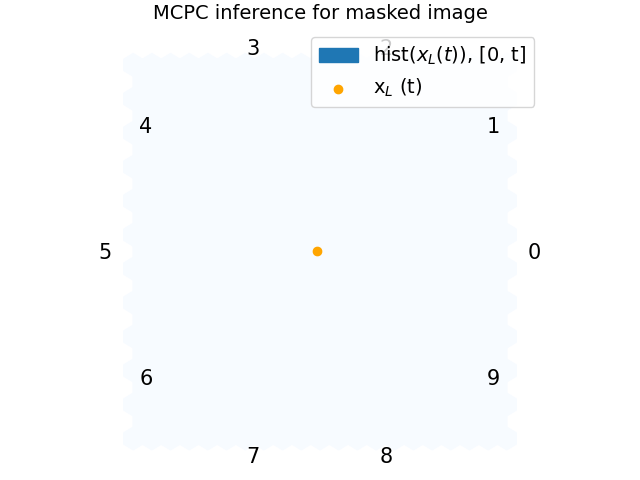

### Supplementary video 3

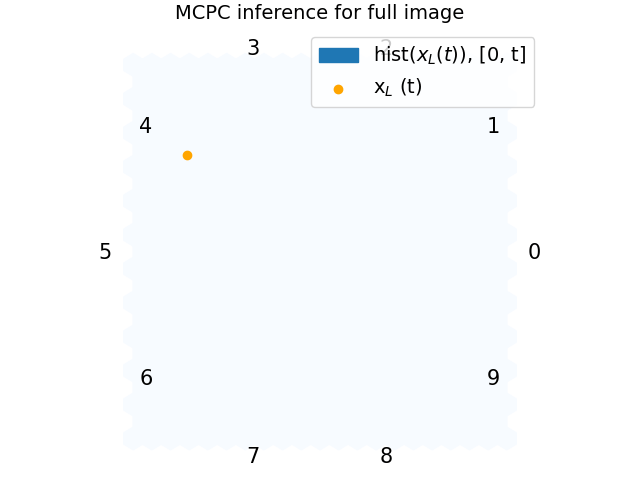

### Supplementary video 4

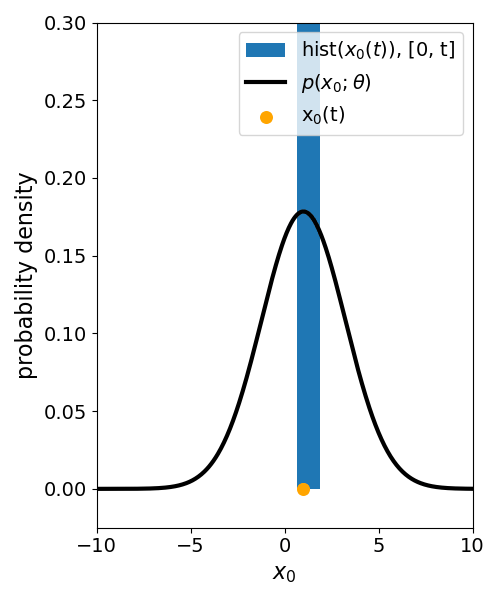

### Supplementary video 5

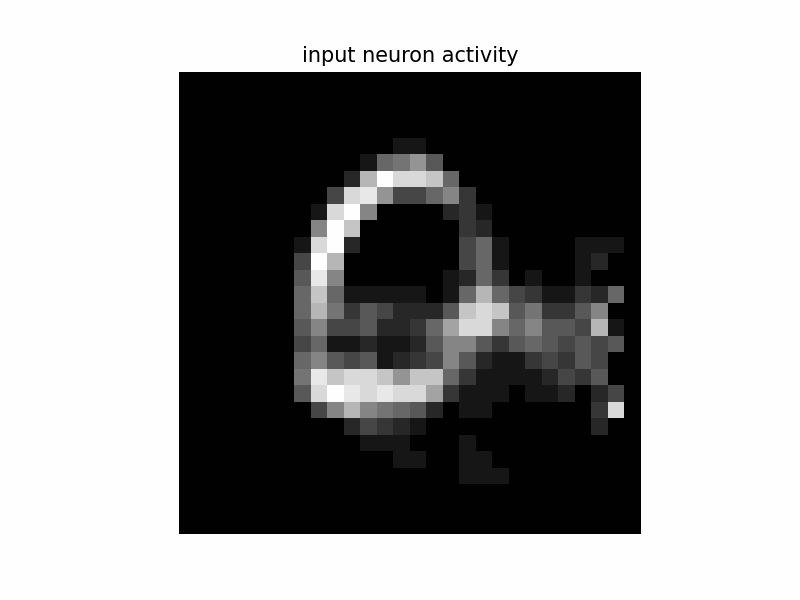
